# A pyridyl-furan series developed from Open Global Health Library blocks red blood cell invasion and protein trafficking in *Plasmodium falciparum* through potential inhibition of the parasite’s PI4KIIIb enzyme

**DOI:** 10.1101/2023.04.25.538349

**Authors:** Dawson B. Ling, William Nguyen, Oliver Looker, Zahra Razook, Kirsty McCann, Alyssa E. Barry, Christian Scheurer, Sergio Wittlin, Hayley E. Bullen, Brendan S. Crabb, Brad E. Sleebs, Paul R. Gilson

## Abstract

With resistance increasing to current antimalarial medicines, there is an urgent need to discover new drug targets and to develop new medicines against these targets. We therefore screened the Open Global Health Library of Merck KGaA, Darmstadt, Germany of 250 compounds against the asexual blood stage of the deadliest malarial parasite *Plasmodium falciparum,* from which eight inhibitors with low micromolar potency were found. Due to its combined potencies against parasite growth and inhibition of red blood cell invasion, the pyridyl-furan compound OGHL250, was prioritised for further optimisation. The potency of the series lead compound (WEHI-518) was improved 250-fold to low nanomolar levels against parasite blood-stage growth. Parasites selected for resistance to a related compound MMV396797, were also resistant to WEHI-518 as well as KDU731, an inhibitor of the phosphatidylinositol kinase PfPI4KIIIB, suggesting this kinase is the target of the pyridyl-furan series. Inhibition of PfPI4KIIIB blocks multiple stages of the parasite’s life cycle and other potent inhibitors are currently under preclinical development. MMV396797-resistant parasites possess an E1316D mutation in PfPKI4IIIB which clusters with known resistance mutations of other inhibitors of the kinase. Building upon earlier studies which showed that PfPI4KIIIB inhibitors block the development of the invasive merozoite parasite stage, we show that members of the pyridyl-furan series also block invasion and/or the conversion of merozoites into ring-stage intracellular parasites through inhibition of protein secretion and export into red blood cells.

---

Malaria is caused by infection with *Plasmodium* parasites and imposes a substantial global burden with there being an estimated 247 million cases and 619,000 deaths in 2021 ^1^. These estimates are an increase of approximately 10% from previous years, which is believed to be largely because of the SARS-CoV-2 pandemic disrupting health services and increasing resistance to currently available antimalarials. *P. falciparum* is the species with which most of the malaria disease burden is associated, and of great concern is the emergence of parasite resistance to frontline artemisinin combination therapies (ACTs) around the globe ^2-6^. This expanding antimalarial resistance highlights an urgent need to discover and develop novel antimalarial medicines that have different mechanisms of action (MoA) to those currently in use, most of which parasites have developed some degree of resistance to. To date, efforts to develop new antimalarials have largely centred on performing phenotypic screens of compound libraries for small molecules that inhibit the growth of the disease-causing asexual blood stage of *P. falciparum*. This has resulted in the screening of several million compounds and the identification of tens of thousands of inhibitory molecules ^7-9^.

To help prioritise which inhibitory compounds to advance, it is important to know the compound’s MoA. It is also crucial to determine whether the compounds target other stages of the parasite’s lifecycle including the pre-erythrocytic liver stage, and the transmission (sexual) stage. Target identification generally involves growing parasites in the presence of the inhibitors of interest until resistance develops, after which complete genome sequencing of the parasite may reveal mutations in the genes encoding the proteins targeted by the inhibitors. Amplification of genes encoding the target proteins can also be selected to improve resistance through increased protein levels. Resistance selection can take many months and does not always yield resistant parasites. Compounds to which resistance cannot be developed *in vitro* are said to be irresistible ^10,11^. Unfortunately, even when resistance can be developed, there is a high likelihood of rediscovering an already well-characterised protein target for which many inhibitors are already known. Common antimalarial targets often detected by these broad screening methods include PfATP4 and DHODH/cytochrome bc1 ^12-14^. Identifying novel compounds targeting these proteins can be partly mitigated by screening compound libraries with parasites already resistant to a common target such as DHODH/cytochrome bc1, or by performing assays for the biological effects known to be caused by common inhibitors such as those that target PfATP4^12,13^.

Another way of finding and prioritising inhibitors with novel MoAs is to perform phenotypic screens for inhibition of biological phenomena such as egress from old red blood cells (RBCs) and invasion of new RBCs. Such screens of the Medicines for Malaria Venture (MMV) Malaria and Pathogen boxes have identified dozens of compounds that inhibit egress and invasion ^15,16^. Novel targets identified by these screens include actin and profilin; essential components of the invasion machinery used by the parasite to enter RBCs ^17^. Egress and invasion inhibitors could be particularly effective compounds since reducing the capacity of invasive merozoite stage parasites to rapidly invade RBCs would leave the merozoites exposed to host immune clearance ^18,19^.

While phenotypic screens for egress and invasion can yield novel targets such as actin and profilin, they are not guaranteed to return solely novel targets ^17^. Indeed, nearly half of the egress and invasion inhibitors identified in recent screens of the MMV Malaria and Pathogen Boxes were also flagged as inhibitors of PfATP4 ^12,13,15,16^. These have later been determined to be specific inhibitors of egress and not invasion ^20^ and serve to dysregulate sodium ion levels and cellular pH, explaining why late schizont-stage parasites are particularly susceptible to the effects of PfATP4 inhibitors ^20,21^. Therefore, while egress and invasion screens can also identify inhibitors to well-studied targets, they can add important additional biological information about the function of targets.

To find inhibitors with novel targets and to reduce the chances of identifying inhibitors of known targets from libraries enriched for antimalarial compounds, we screened the Open Global Health Library (OGHL) of Merck KGaA, Darmstadt, Germany containing 250 compounds ^22^. A screen of this library for compounds that reduce the growth of *P. falciparum* parasites identified several inhibitors of low micromolar potency or less. Subsequent egress and invasion inhibition assays indicated half of these compounds were also invasion inhibitors with the pyridyl-furan compound OGHL250 being among the most potent. OGHL250 was selected for further improvement by medicinal chemistry and the potency of its analogues was greatly increased against parasite growth and invasion. As the pyridyl-furan analogues were structurally related to the compound MMV396797 (M-797), we likewise assayed it against invasion and found it was inhibitory too. Recently M-797 was shown to block parasite protein secretion and export^23^ and so we evaluated the pyridyl-furan analogues and found they too inhibited protein secretion and export. Furthermore, the M-797-resistant parasites were also resistant to the pyridyl-furan analogues and an inhibitor of the protein trafficking enzyme phosphatidylinositol 4-kinase (PfPI4KIIIB), suggesting it as the common target. Given this, we believe the pyridyl-furan series of analogues blocks protein trafficking and invasion by inhibiting PfPI4KIIIB.

## RESULTS

### Screening of the OGHL for inhibitors of *P. falciparum* growth identified eight inhibitory compounds

To screen the OGHL library for parasite growth inhibition, *P. falciparum* parasites transfected with a nanoluciferase (Nluc) reporter protein exported into the RBC compartment were used as these parasites were required for our downstream egress and invasion assays ^15,24^. The OGHL library was diluted to 2 µM in RPMI media and in it were grown young ring-stage Nluc parasites for 72 hours during which they passed through 1.5 cell cycles. The biomass of the older trophozoite/schizont-stage parasites was assayed by measuring parasite lactate dehydrogenase (LDH) activity which indicated seven compounds inhibited growth by >50% relative to a DMSO drug vehicle control (Figure 1A, Table S1). An eighth compound L15, was slightly above the 50% cut-off and was retained as it was below the cut-off for Nluc inhibition (see next). In parallel, assays measuring Nluc activity were also performed and indicated nine OGHL compounds inhibited growth by >50% of which eight compounds were common with those identified by the LDH assay (Figure 1B, Table S1).

**Figure 1.**
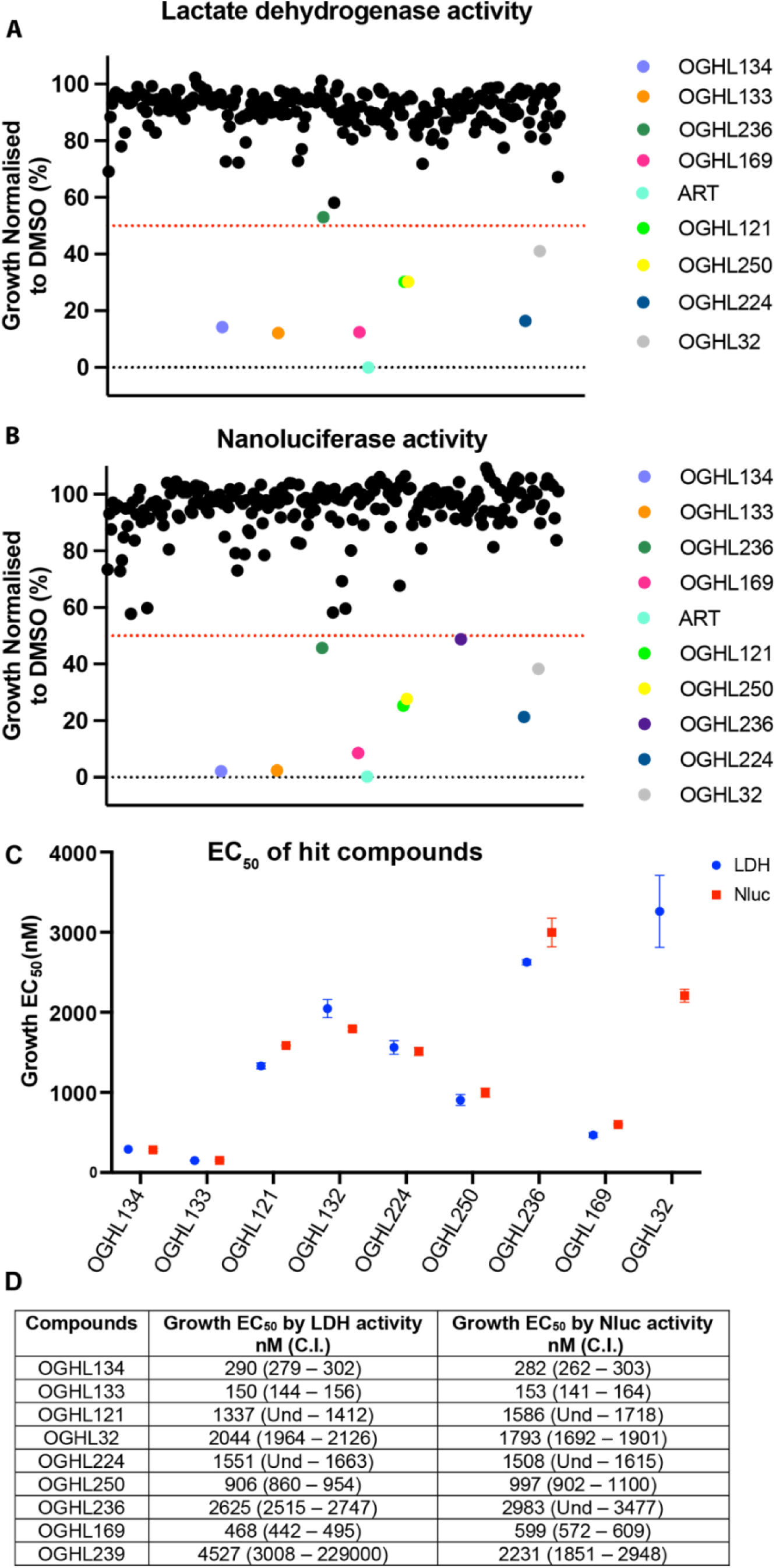
Using *P. falciparum* parasites expressing an exported nanoluciferase (Nluc) reporter, several Open Global Health Library (OGHL) compounds at 2 µM demonstrated ≥ 50% blood-stage growth inhibition after a 72-hour growth period. (A) Parasite growth after drug treatment was measured using lactate dehydrogenase activity (LDH) and seven compounds inhibited ≥ 50%. (B) Following measurement of Nluc activity, nine OGHL compounds were found to inhibit growth by ≥50%. All values were normalised to vehicle control (0.1% DMSO) and each dot represents the mean of a compound from three technical replicates. Dotted lines indicate cut-off values of 50% for growth inhibition. Artemisinin (ART) was used as a positive control at 80 nM. (C and D) Graphical and numerical EC_50s_ of OGHL hit compounds after 72-hours growth in serially diluted compounds. Growth was measured by LDH (blue) and Nluc activity (red).

Supplementary compound of the screening hits and their structures were provided by Merck KGaA Darmstadt, Germany to enable downstream experiments (Figure S1)^22^. To derive the EC_50_ for growth, the nine new OGHL hit compounds were serially diluted and Nluc parasites were grown for 72 hours. EC_50_ values were calculated for both LDH and Nluc assays, and they were in close agreement, except for OGHL239 (Figure 1C and 1D and Figure S1 and S2). A counter screen was therefore performed using freshly prepared cell lysates of the Nluc parasites. In this screen compound OGHL239 was observed to strongly inhibit Nluc activity leaving eight inhibitors with about 50% growth inhibitory activity (and no specific activity against the Nluc) to be further studied (Figure S3).

### None of the OGHL growth inhibitors affected egress of *P. falciparum* parasites

To measure normal and premature egress of merozoite-stage parasites and their invasion of new RBCs, we employed a previously developed assay (workflow is shown in Figure 2A) ^15^. One of the key events which helps trigger the egress of merozoites from schizonts is the stimulation of protein kinase G (PKG) activity through an increase in cyclic GMP (cGMP) levels ^25,26^. cGMP-activated PKG triggers a signalling cascade that results in the merozoites releasing proteases and other molecules that help breakdown the RBC so the merozoites can egress and invade new RBCs ^27^. To prevent premature activation of PKG, phosphodiesterases (PDEs) hydrolyse cGMP, reducing levels of the cyclic nucleotide. PDE inhibitors such as zaprinast lead to a rapid increase in cGMP levels triggering premature activation of PKG and subsequent accelerated egress of immature, poorly invasive merozoites ^25,26^. Since the OGHL library contains many experimental human PDE inhibitors, we assayed Nluc parasites for accelerated egress 30 minutes after adding the OGHL compounds at 10x EC_50_ for growth ^22^. After a 30-minute premature egress window, media containing Nluc released from the infected RBCs was harvested and supplemented with NanoGlo substrate with the bioluminescence measured being proportional to the amount of egress that had occurred ^15,26^. Relative to egress activator zaprinast, none of the OGHL compounds appeared to accelerate egress suggesting they were not inhibitors of *P. falciparum* PDEs (Figure 2B).

**Figure 2.**
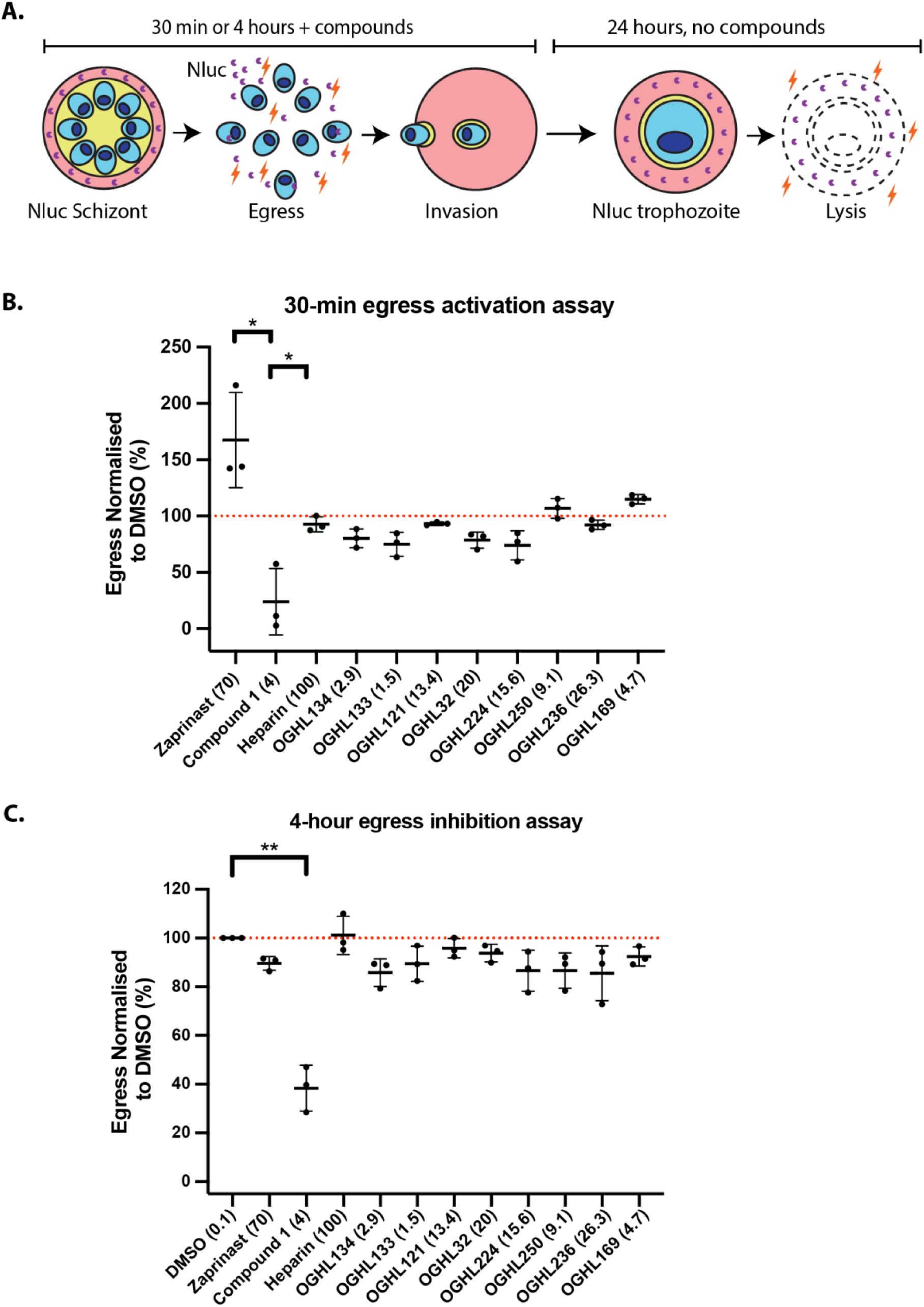
OGHL hit compounds did not activate egress in the short term (30 mins) or inhibit in the longer term (4 hours). (A) Workflow summary of how the egress and invasion assay works. (B) Late-stage schizonts were treated with the OGHL compounds for 30 minutes, and the growth medium was collected to measure Nluc bioluminescence as a marker of egress. None of the OGHL compounds appeared to activate egress. Zaprinast was used as positive control egress stimulator (167% compared with 0.1% DMSO control normalised to 100%), while C1 was an egress inhibitor (24% of DMSO control). Heparin was an invasion inhibitor and as expected had no effect on egress as expected. OGHL compounds were tested at 10x EC_50_ of growth and all concentrations specified in brackets are in µM, except for heparin (in μg/mL). (C) Over a four-hour window, none of the OGHL compounds appeared to significantly inhibit egress, compared to the DMSO control. C1 efficiently blocked normal egress to 38% compared to the 0.1% DMSO control, normalised to 100%. Error bars represent the standard deviation of three biological replicates, each with three technical replicates. Dotted line indicates cut-off value of 100% for egress activity. Statistical analyses were performed on GraphPad Prism 9 with Welch’s t-test between DMSO and C1. **P < 0.01, *P < 0.05. No bar indicates not significant.

To examine the ability of the OGHL inhibitors to inhibit ‘normal’ egress, we next assayed the OGHL inhibitors at 10x EC_50_ (for growth) for their capacity to inhibit egress over a 4-hour window in which most merozoites will have naturally egressed using the same Nluc release assay as used for the premature egress analysis. Briefly, transgenic late schizont-stage parasites exporting Nluc into the RBC compartment were allowed to egress and invade in the presence of 10x EC_50_ (for growth) of each of the eight OGHL compounds. After the 4-hour egress and invasion window, media containing Nluc released from the infected RBCs was harvested and supplemented with NanoGlo substrate as before (Figure 2A). Relative to the control DMSO, the OGHL compounds did not significantly inhibit egress (Figure 2C). The control PKG inhibitor Compound 1 (C1) did however substantially reduce egress as anticipated 15.

### Half of the OGHL inhibitors reduced *P. falciparum* invasion of RBCs

Following the 4-hour egress assay, the parasites were treated to remove both the unruptured schizonts and the OGHL compounds leaving the newly invaded ring-stage parasites to continue to grow. The young parasites were grown for a further 24 hours in the absence of OGHL compound, whereupon the older trophozoite-stage parasites were lysed and the Nluc activity quantified as an indicator of invasion success ^15^. Compared to the DMSO control, compounds OGHL133 and OGHL224 did not inhibit invasion at all (Figure 3). In contrast, compounds OGHL32, OGHL250, OGHL236 and OGHL169 strongly inhibited invasion below the 50% cut off, with compounds OGHL134 and OGHL121 being borderline. Overall, OGHL250 had the strongest combination of low EC_50_ for invasion and growth, therefore we decided to optimise this compound for greater potency against *P. falciparum,* followed by efforts to derive the compound’s MoA.

**Figure 3.**
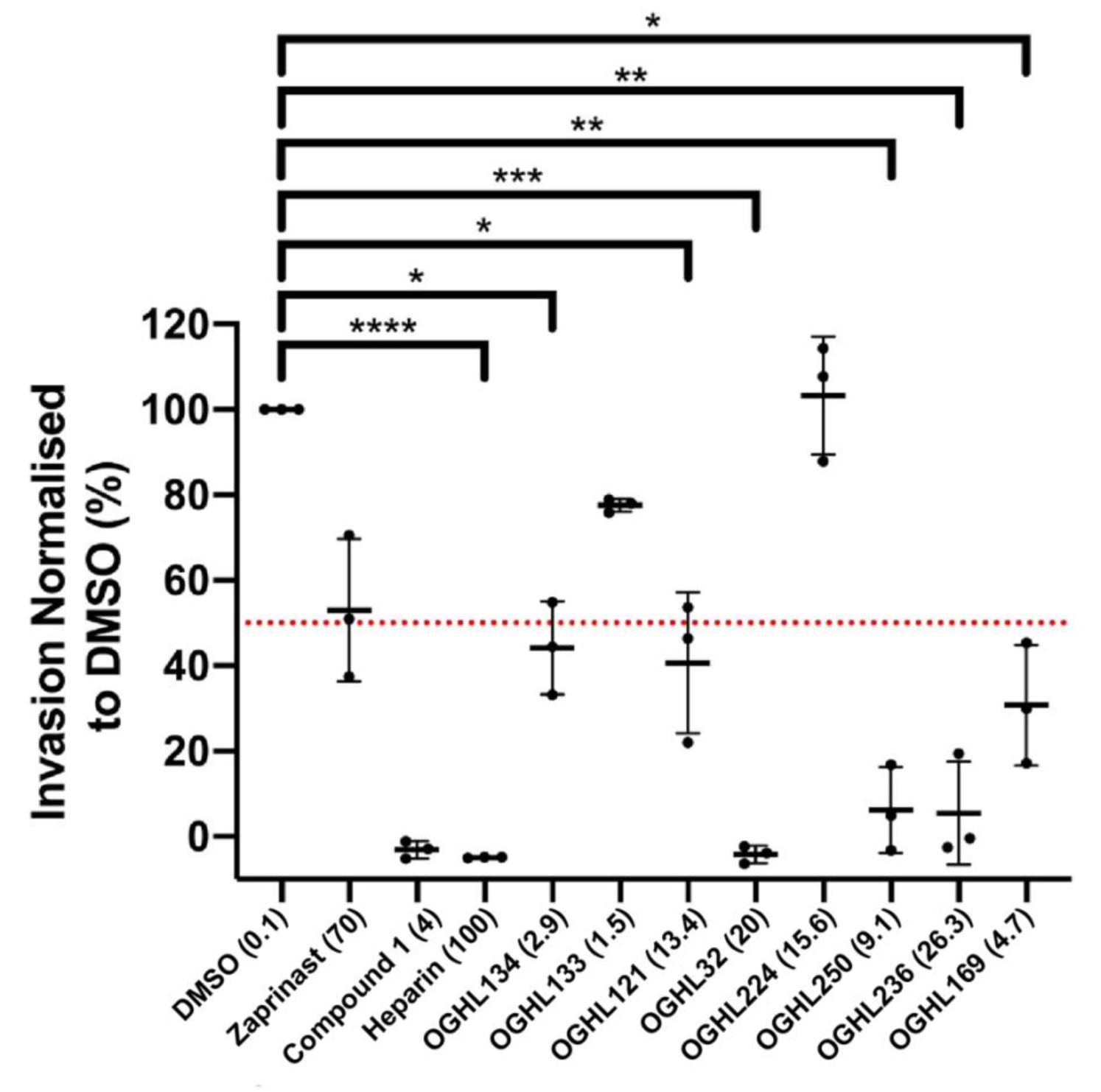
Six of eight OGHL compounds demonstrate varying degrees of invasion inhibition when used at 10x EC_50_ for growth. When normalised to 0.1% DMSO, the OGHL invasion inhibitors displayed with almost complete invasion inhibition for OGHL32 and control compound, heparin, <10% invasion for OGHL250 and OGHL236 and <50% inhibition OGHL134, OGHL121 and OGHL169. All OGHL compounds are in μM, except for heparin (in μg/mL) and DMSO (in %). Error bars represent the standard deviation of three biological replicates, each with three technical replicates. Dotted line indicates cut-off value of 50% for invasion activity. Statistical analyses were performed on GraphPad Prism 9 with Welch’s t-test between DMSO and control compound, heparin, and OGHL compounds. *P < 0.05, **P < 0.01, ***P < 0.001, ***P < 0.0001. No bar indicates not significant.

### Optimization of pyridyl-furan parasite activity

The optimization of the anti-parasitic activity of the pyridyl-furan **OGHL250** hit chemotype began with relocating the endocyclic nitrogen on the isoquinoline ring to the 6-position (**1**) (Figure 4). This modification resulted in a 10-fold improvement in parasite activity (EC_50_ 0.33 µM) while improving the selectivity window against human HepG2 cells (CC_50_ 10 µM). The addition of basic functionality by way of an *N*-methyl piperazine (**2**) group onto the carboxamide resulted in a 6-fold improvement in parasite activity (EC_50_ 0.89 µM) relative to **OGHL250**. These two modifications were then combined in the analogue **3** and resulted in a 2-fold improvement in parasite activity (EC_50_ 0.16 µM) relative to **1**. Removal of the methoxy substitution from the benzamide aryl ring (**WEHI-518**) led to a further 7-fold improvement in activity (EC_50_ 0.022 µM).

**Figure 4.**
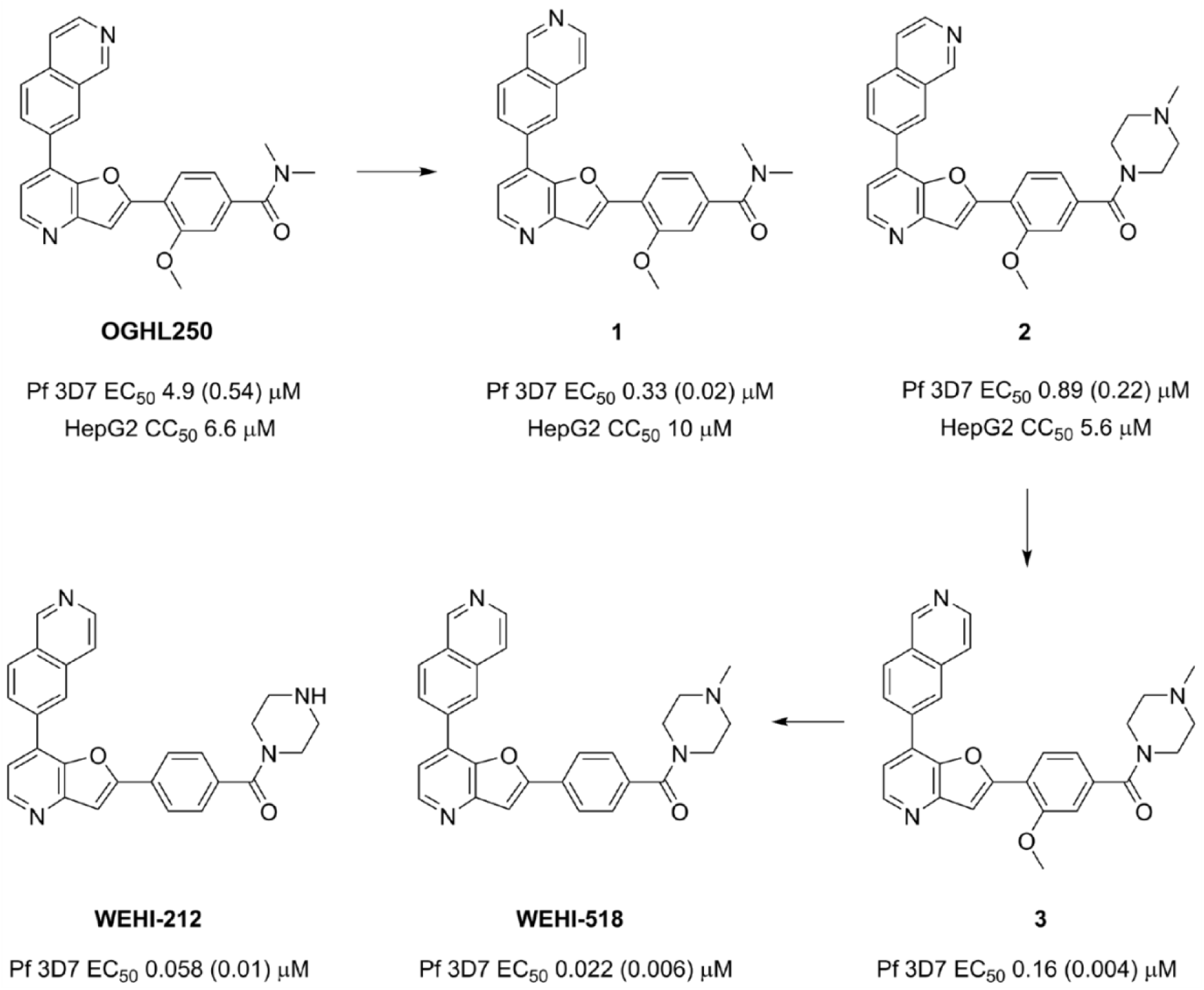
Synthesis of optimized analogues of OGHL250showing EC_50_s against blood-stage *P. falciparum* parasites and CC_50_ against human HepG2 cells. EC_50_ data for *P. falciparum* are averages of three biological replicates and SD in brackets. HepG2 toxicity data are shown for the first three compounds with n=1.

The metabolic stability of **WEHI-518** was determined by incubation in mouse liver microsomes. It showed that **WEHI-518** has low metabolic stability (CL_int_ 301 μL/min/mg) (Table S2). It was speculated that the methyl substitution on the piperazine of **WEHI-518** was susceptible to metabolic N-demethylation. **WEHI-212** without the *N*-methyl group was synthesized and was shown to retain parasite activity (EC_50_ 0.058 µM) and improved metabolic stability (CL_int_ 180 μL/min/mg). This indicated that there are other sites of metabolism in **WEHI-518** and **WEHI-212** and these are likely to include cytochrome P450-mediated oxidation of one or both pyridyl nitrogens and oxidation of the aryl group. Determination of kinetic aqueous solubility established **WEHI-518** possessed high aqueous solubility (160-320 µM) while **WEHI-212** was modest (20 - 40 µM).

In summary of the optimization of the pyridyl-furan chemotype, several key modifications installed in **WEHI-518** were made that led to a 250-fold improvement in parasite activity compared to the hit **OGHL250**. These modifications also inadvertently led to the activity of **WEHI-518** against human kinases (IC_50_ 0.22 to >10 µM) (Table S3) compared to potent human kinase activity observed by the hit chemotype ^28,29^. As a consequence of moderately tuning out the kinase activity, the HepG2 selectivity window was also improved. The physicochemical properties installed ensured **WEHI-518** has high aqueous solubility, however, further optimization is required to improve the metabolic stability to a level suitable for *in vivo* efficacy studies in mouse malaria models.

### Pyridyl-furan analogues of OGHL250 block parasite invasion but not egress

Following development of the pyridyl-furan series, we repeated the 4-hour egress and invasion assay as described in Figure 2 with the three lead compounds (**OGHL250**, **WEHI-518** and analogue **3**[renamed **Compound 3** or **C3**]), but this time challenged the parasites with a dilution series of the compounds to generate EC_50_s (Figure 5A–C). As anticipated, **WEHI-518** and **C3** like **OGHL250**, did not inhibit parasite egress. In contrast, all the pyridyl-furan compounds blocked invasion over 4 hours in the same general order of potency as in the 72-hour growth assay in which we included the additional antimalarial resistant Dd2 strain (Figure 5E). Generally, the pyridyl-furan compounds inhibited growth more strongly than invasion suggesting they could target multiple stages of the asexual blood stage cell cycle (Figure 5B).

**Figure 5.**
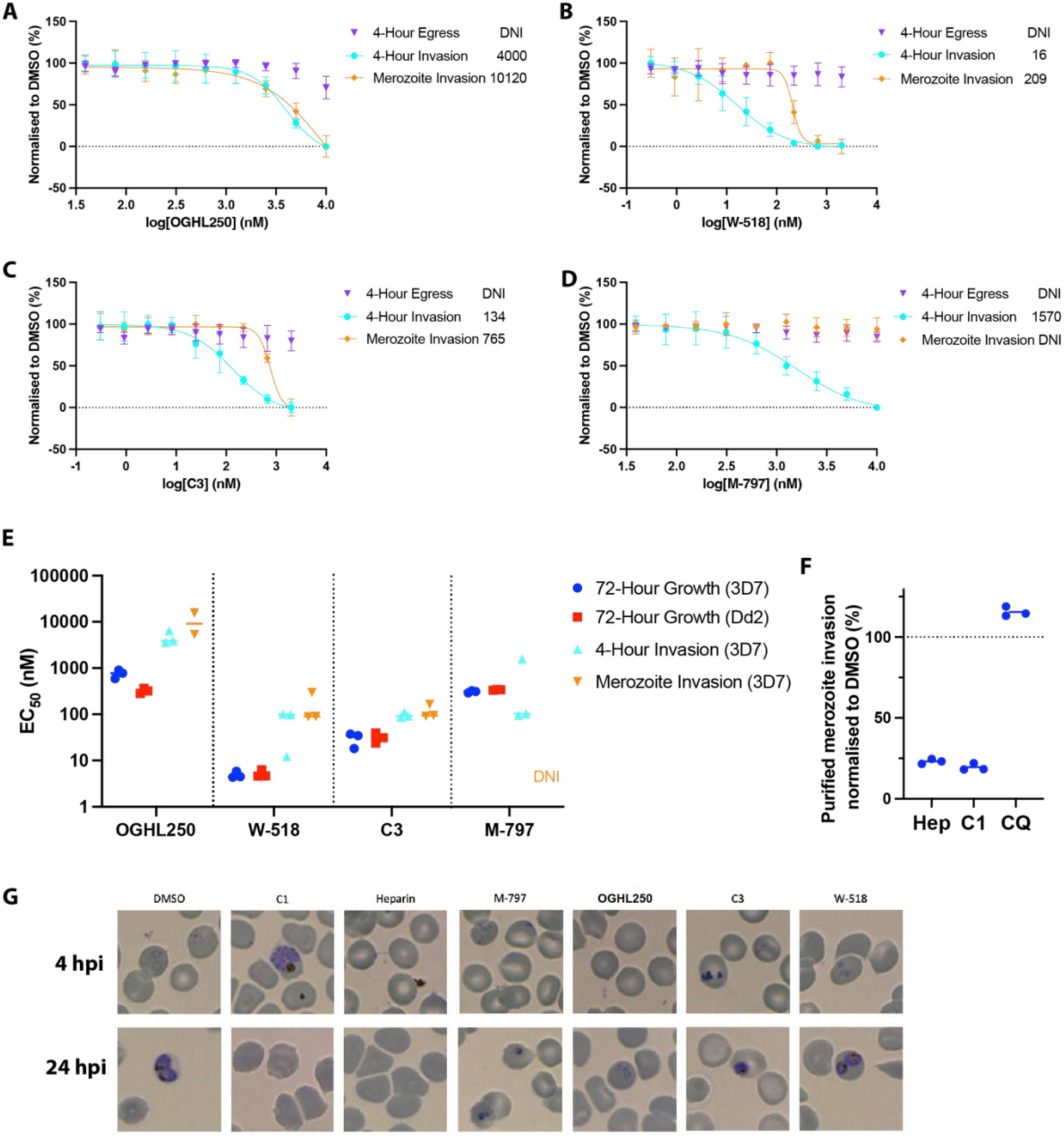
M-797 and pyridyl-furan compounds block invasion but not egress during a 4-hour assay. (A-D) Dose response curves for 4-hour egress and invasion assays for **OGHL250**, **WEHI-518**, **C3** and M-797 respectively, indicate invasion but not egress was blocked. Invasion inhibition of purified merozoites indicates all the compounds except for M-797 block RBC invasion. EC_50_ values (nM) for each type of assay are indicated on the right of each graph. Error bars indicate SD of 3 biological replicates. (E) EC_50_ values of 3D7 and Dd2 parasites treated with the compounds indicated for 72-hours and EC_50_s for the 4-hour invasion and purified merozoite invasion assays with Nluc parasites are shown for 2-3 biological replicates. (F) The percentage of invasion of purified merozoites relative to 0.1% DMSO are shown for heparin (100 µg/mL), C1 (4 µM) and CQ (chloroquine, 75 nM) for 3 biological replicates. (G) Giemsa-stained thin blood smears of *P. falciparum* parasites after the 4-hour egress/invasion assay indicate the pyridyl-furan compounds do not specifically block merozoite invasion as some ring-stage parasites are present. The parasite cultures were treated with DMSO (0.12%), compound 1 (C1; 4 µM), heparin (100 µg/mL), M-797 (4.71 µM), **OGHL250** (12 µM), **WEHI-518** (**W-518**, 0.05 µM) and **C3** (0.4 µM). Blood smears made after the 4-hour drug treatment window (4-hour post-invasion [hpi]) and after the 24-hpi show the ring-stage parasites failed to thrive after invasion after treatment with the OGHL250 series and M-797. DNI = did not inhibit.

Since we have recently shown that M-797 blocked protein trafficking in parasites and is structurally related to **OGHL250** (Figure S4) we decided to determine if M-797 could block egress and invasion ^23^. As RBC invasion requires the trafficking of many invasion proteins to specialised secretory organelles in the merozoite it was anticipated that M-797 might block invasion ^30^. Like pyridyl-furans, M-797 did not block egress but did block invasion over 4 hours with a similar EC_50_ as 72-h growth inhibition (Figure 5D).

To determine if the pyridyl-furan compounds were disturbing the development of viable merozoites or if they were directly blocking specific invasion processes, schizonts containing merozoites were allowed to mature in the absence of inhibitors. The mature merozoites were then purified and allowed to invade RBCs in the presence of the compounds. After 30 mins, the compounds were removed, and the newly invaded parasites were grown for 24-hours to enable quantification of their Nluc levels. We found that **OGHL250**, **WEHI-518** and **C3** inhibited invasion with EC_50_ concentrations higher those found in the 4-hour egress and invasion assay (Figure 5A–C and 5E), indicating that the compounds were not only targeting merozoite development, but were also directly blocking invasion and/or early-stage ring conversion. M-797 on the other hand, did not inhibit merozoite invasion suggesting its target was required for merozoite development rather than invasion and/or conversion (Figure 5D and E). For merozoite invasion we also included an invasion-blocking heparin control which performed as anticipated (Figure 5F)^15^. The antimalarial chloroquine had no effect on invasion and the protein kinase G inhibitor C1, also directly inhibited merozoite invasion which has been noted before with another PKG inhibitor (Figure 5A)^15,31^.

To gain further insights into the compounds’ inhibitory mechanisms, Giemsa-stained blood smears were examined after completion of the 4-hour egress and invasion assay (Figure 5G). In DMSO-treated cultures, many healthy, rounded ring-stage parasites were observed and as anticipated, C1 treatment blocked merozoite egress and heparin blocked invasion (Figure 5G). For M-797, **OGHL250**, **C3** and **WEHI-518** small, misshapen ring-stage parasites were observed suggesting some invasion had occurred but that conversion into healthy ring-stage parasites was inhibited. By 20 hours later, the M-797, **OGHL250**, **C3** and **WEHI-518**-treated parasites had failed to thrive and were noticeably smaller (Figure 5G).

### The Pyridyl-furan analogues but not OGHL250 block protein secretion and export in parasites

As M-797 had been previously shown to block parasite protein secretion and export into the RBC compartment^23^, we next explored the effects of the pyridyl-furans on protein secretion and export. We used a recently developed parasite protein trafficking inhibition assay ^23^ designed to detect inhibitors of protein trafficking both within the parasite and at the specialised parasitophorous vacuole membrane (PVM, the membranous sac containing the parasite within the infected RBC) (Figure 6A). Specifically, we treated parasites with a selection of protein trafficking inhibitors including brefeldin A, torin 2 and M-797 ^23,32,33^ (Figure S4). In this assay, young Nluc trophozoite-stage parasites were treated with compounds for 5 hours to block secretion into the parasitophorous vacuole (PV) and export into the RBC compartment after which they were differentially lysed in reagents designed to release the contents of the RBC only, RBC and PV or all compartments of the infected RBC (Figure 6A). Lysis of these compartments permitted Nluc to gain access to its substrate and produce a bioluminescent signal proportional to Nluc trapping within that compartment ^23^. Relative to the DMSO control, brefeldin A, torin 2 and M-797 displayed significantly reduced export into the RBC compartment and therefore greater accumulation in the PV (secreted) and parasite (retained) compartments, as expected for protein trafficking inhibitors (Figure 6B). While OGHL250 did not significantly reduce export of the protein reporter into the RBC compartment, pyridyl-furan analogues **WEHI-518** and **C3,** as well as putative protein trafficking inhibitor KDU731 ^34^ behaved similarly to the control export blocking compounds and showed reduced protein export into the RBC compartment with stronger protein trapping at the PV and parasite compartments (Figures 6B and S4).

**Figure 6.**
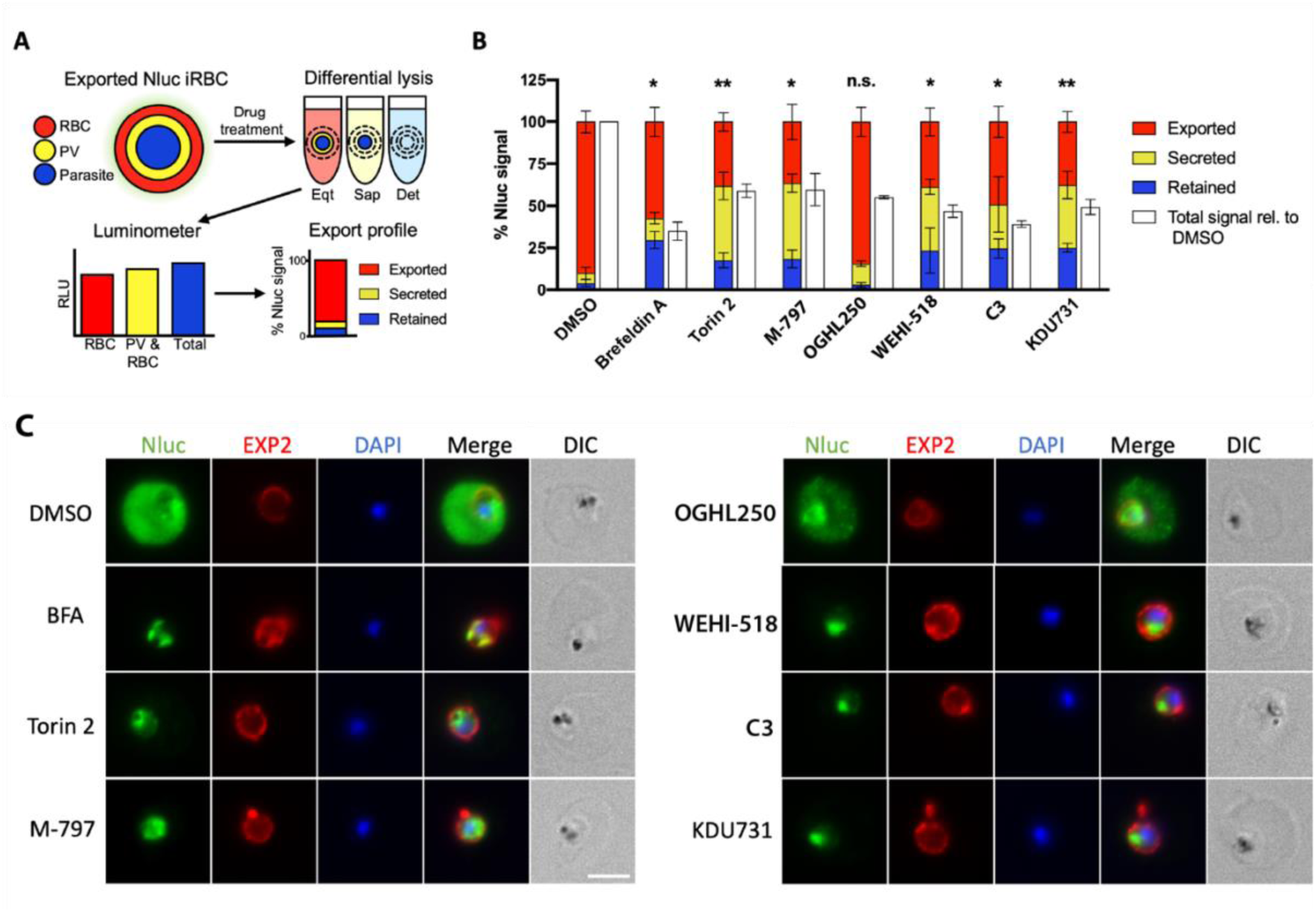
PfPI4KIIIB inhibitors reduce protein secretion and export in *P. falciparum*. (A) Workflow diagram illustrating how protein export screen works to measure retention of Nluc reporter in the different compartments of infected RBCs. (B) The percentages of Nluc reporter retained in the parasite, secreted into the PV or exported into the RBC compartments after 5 h of compound treatment are indicated. Error bars indicate the standard deviations of three biological replicates of three technical replicates. Compound used are; DMSO (0.1%), Brefeldin A (5 µg/mL), Torin 2 (10 nM), M-797 (4.8 µM), **OGHL250** (3.1 µM), **WEHI-518** (200 nM), **C3** (1.2 µM) and KDU731 (300 nM). The total Nluc signal relative to DMSO control are indicated by white bars. Welch’s T tests were performed comparing the %Exported signal for each compound versus DMSO using GraphPad PRISM. *p <0.05, **p<0.01. (C) Fluorescence microscopy images of parasite infected RBCs treated with the same inhibitory compounds used in B. The cells were probed with rabbit anti-Nluc and an EXP2 mouse monoclonal to locate the Hyp1-Nluc reporter protein and the PVM, respectively. Nuclei were stained with DAPI and the space bar indicates 4 µm.

### OGHL250 analogues act like PfPI4KIIIB inhibitors

To identify the protein target and mechanism of action of the pyridyl-furan compounds, we attempted to select for genetic resistance to OGHL250 through repeated rounds of treatment and recovery against engineered ‘hypermutator’ Dd2 parasites that have a higher background error prone DNA replication and are hereafter called Parental parasites ^35,36^. After several rounds of selection, the DMSO-treated Parental parasites and OGHL250-challenged parasites were subjected to a 72-hour growth assay against a dilution series of OGHL250. However, the EC_50_ was little changed, indicating no resistance could be selected for (data not shown).

In parallel, we selected for resistance to M-797 using the mutator Parental parasites as we were interested in the protein trafficking target of this compound. The Parental parasites were cycled three times on and off M-797 and resistance to the compound was measured via a 72-hour growth assay where four lines with about 16-fold resistance were recovered (Figure 7 and S5A). The M-797-resistant parasites lines were subsequently tested against torin 2 and were not resistant, but they were 8 to 13-fold more resistant to **WEHI-581**, **C3** and KDU731 than the Parental control parasites (Figure 7 and Figure S5B-E). However, the M-797 resistant parasites were not resistant to OGHL250, which was surprising as OGHL250 did not inhibit protein trafficking (Figure 7 and S5F). Overall, these data suggest that the pyridyl-furan analogues, M-797 and KDU731 have a shared target.

**Figure 7.**
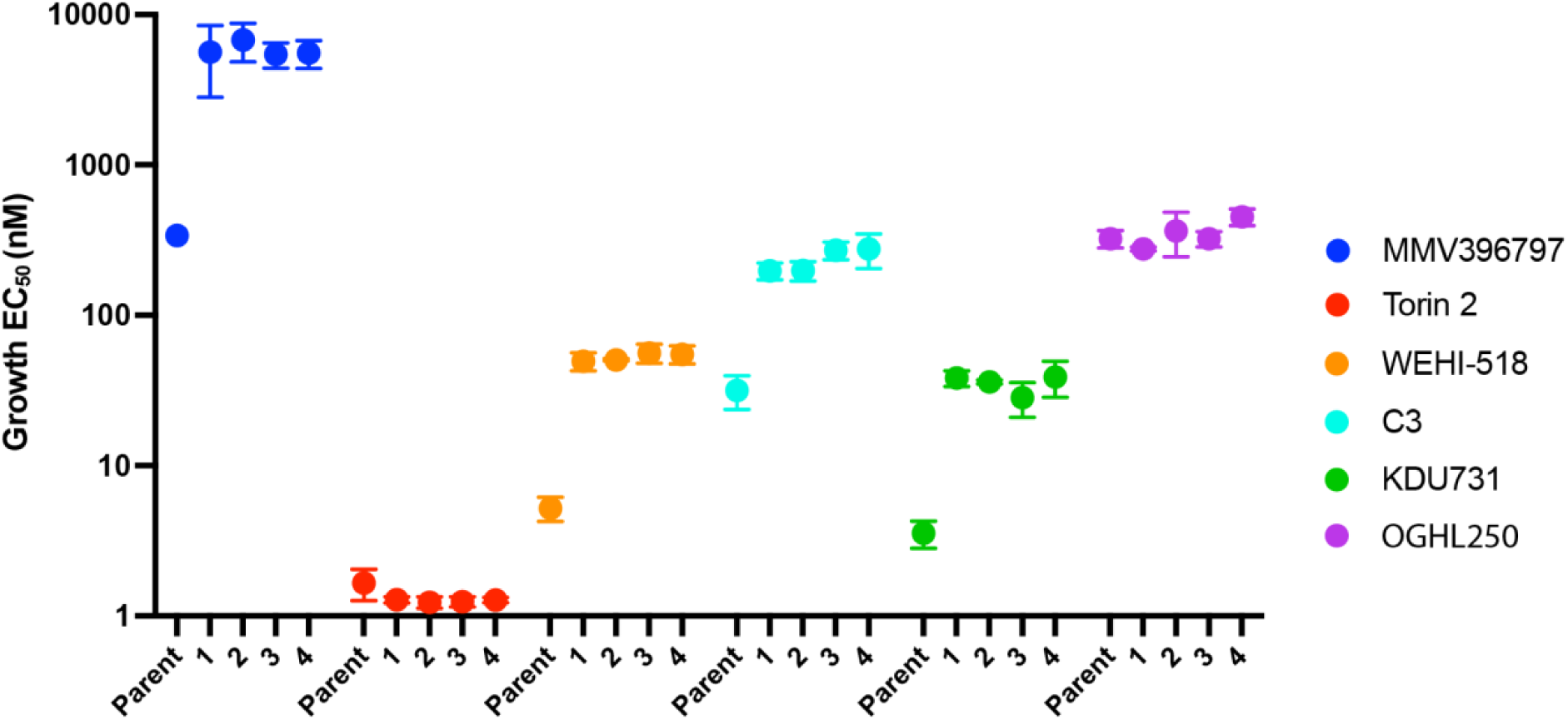
M-797-resistant parasites are resistant to some but not all PfPI4KIIIB inhibitors. Four parasite lines selected for resistance to M-797 as well as the non-resistant Parent parasites were grown for 72 hours in serially diluted inhibitors and EC_50_s for growth inhibition were calculated. The EC_50_s for three biological replicates each of three technical replicates are plotted with bars indicating standard deviations. Y-axis scale represents antilog of drug concentrations in nM.

### Parasites resistant to M-797 have a mutation in their PfPI4KIIIB gene

As torin 2 analogue, NCATS-SM3710, is a known inhibitor of PfPI4KII*B* (PF3D7_0509800) and KDU731 inhibits the PI4K enzyme of *Cryptosporidium spp*. we predicted that M-797, **WEHI-518** and **C3** might also be PI4KIII*B* inhibitors ^34,37^. PfPI4KIIIB (PF3D7_0509800) is 1559 aa in size and the last three-quarters of the protein is predicted to comprise a phosphatidylinositol 4-kinase beta domain in which resistance mutations to other PI4K inhibitors have been found (Figure 8A). As a first step to address this, M-797, **OGHL250** and **WEHI-518** were assayed with the previously described ^3^H-hypoxanthine incorporation assay against a parasite line resistant to MMV390048 that has a S743T mutation in PI4KIIIB ^38^. Compared to the parental Dd2 parasites, the S743T parasites were two-fold and 4-fold more resistant to M-797 and WEHI-518, respectively but not OGHL250 (Table S4).

**Figure 8.**
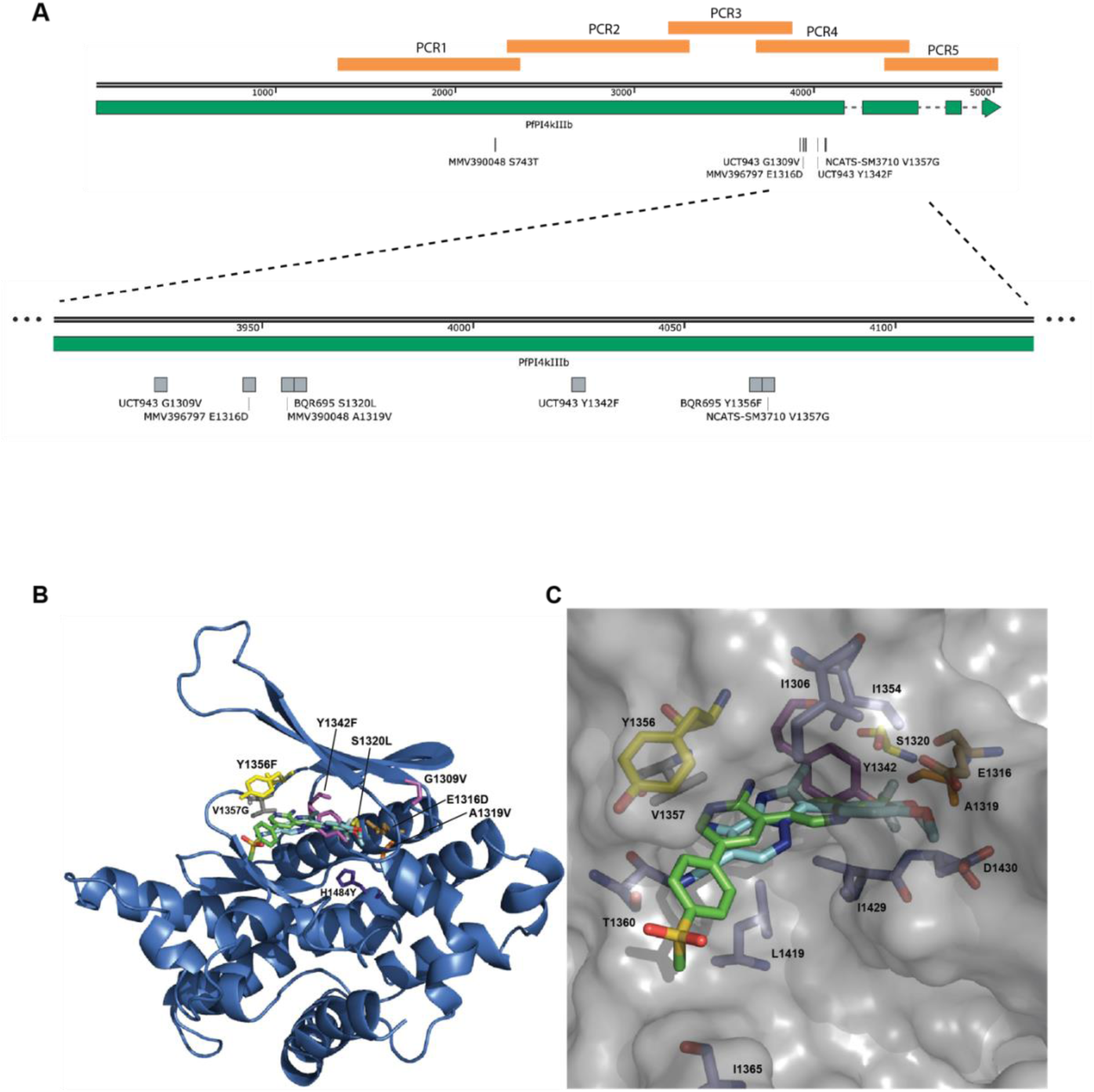
M-797 resistance mutation in *pfpi4kiiib* gene clusters with resistance mutations to other inhibitory compounds. (A, Top) Diagram of whole *pfpi4kiiib* gene showing exons (green) and introns (dashed line), location of PCR amplifed gene fragments and all known resistance mutations. Image made with SnapGene. (b, bottom) Region of *pfpi4kiiib* containing cluster of resistance mutations has been expanded to details. (B) Homology model of the Pf catalytic domain of PI4K (PF3D7_0509800) was built using the PDB 3ENE as previously described by McNamara et al. ^39^. MMV390048 (green) was docked using CLC Drug Discovery workbench. M-797 (cyan) was overlayed using the X-ray structure of this chemotype bound to human PI4KIIIB (PDB 4WAG) ^40^. Mutations conferring drug resistance are shown: E1316D (gold) for M-797; A1319V (orange) for MMV390048; G1309V and Y1342F (magenta) for UCT943; S1320L, Y1356F (yellow) for BQR695; V1357G (grey) for NCATS-SM3710; H1484Y (purple) for KAI407. (C) Surface presentation showing a close-up of the ligand binding site with nearby amino acids shown.

To determine if the M-797-resistant parasites have a mutation in their *pfpi4kiiib* gene, genomic DNA was extracted from the four M-797 resistant lines as well as the Parental parasites. The PI4K domain of PF3D7_0509800 was amplified as a series of 5 overlapping fragments by PCR from one resistant line and the Parental line and compared to *pi4kiiib* of Dd2 (Figure 8A). In the resistant line but not the Parental line, a GAA-GAC mutation was found that encoded an E1316D mutation in PCR4 fragment near to other known drug resistance mutations (Figure 8A). PCR4 fragments from two clones of each of the other three resistant lines were also sequenced, and two of the other lines also had the same GAA-GAC mutation. The remaining line contained a different GAA-GAT mutation at the same position which also encoded the same E1316D mutation. The occurrence of different nucleotide mutations that were independently derived but produced the same amino acid change made us confident that resistance to M-797 was due to this mutation.

To determine if the various resistance mutations mapped closely to each other in the predicted 3D structure a homology model of the *P. falciparum* catalytic domain of PI4KIIIB was built using the PDB 3ENE as previously described by McNamara et al. ^39^. M-797 was predicted to overlap with MMV390048 when overlayed using the X-ray structure of this chemotype bound to human PI4KIIIB (Figure 8B and C) ^40^. Mutations conferring drug resistance are shown: E1316D for M-797, A1319V for MMV390048, G1309V and Y1342F for UCT943, S1320L and Y1356F for BQR695, V1357G for NCATS-SM3710 and H1484Y for KAI407 all appear to be near each other. Interestingly, the MMV390048 S743T resistance mutation is located some distance upstream of the other mutations. These parasites are however, known to be resistant to UCT943 which in turn has its own resistance mutation at Y1342F located near the M-797 resistance mutation E1316D. Alpha-fold structures of PfPI4KIIIB indicate that S743 could project towards the predicted binding sites of the inhibitors providing a mechanism for the resistance of S743T parasites against MMV390048 and other inhibitors (Figure S6)^41-43^.

## DISCUSSION

Here we screened the 250 compound OGHL library of Merck KGaA Darmstadt, Germany for novel inhibitors of merozoite egress and invasion by first performing a 72-h growth inhibitory screen against the asexual blood stage of *P. falciparum*. This resulted in eight compounds being identified with low micromolar potency. This subset was further screened and although none were observed to inhibit egress, four strongly inhibited parasite invasion of human RBCs. Compound OGHL250 was selected for further optimisation as it had a strong combination of growth and invasion inhibition. A 250-fold improvement in growth inhibition was achieved with the series-leading **WEHI-518** compound, but high metabolic turnover in liver microsomes precluded further evaluation in the *P. berghei* mouse model of malaria. **WEHI-212** was subsequently developed to limit metabolic turnover and although this was reduced, it was not significant enough to consider evaluation in *P. berghei*. **WEHI-518** has some mild inhibitory activity against human IKKe and Syk kinases and this will need to be reduced along with improvements in the metabolic stability of the series before further evaluation in mouse parasite models can occur.

Attempts to identify the target of **OGHL250** through selection of parasite resistance to the compound were unsuccessful. **OGHL250** is a known inhibitor of human spleen tyrosine kinase (Syk) for which inhibitory compounds have been developed to treat several diseases ^44^. Given **OGHL250’s** modest potency against parasite growth and poor inhibition of protein export, the compound could have multiple targets in *P. falciparum* including one or more of the parasite’s many kinases^45^. Activity against multiple parasite proteins would decrease the chances of recovering multiple resistance mutations in a single parasite which may have been why resistance to **OGHL250** was not recovered.

Resistance was however recovered to the protein export blocking M-797 compound whose core structurally resembles that of the OGHL250 series prompting us to evaluate the resistance of these parasites to **WEHI-518** and **C3**. As the M-797-resistant parasites were also resistant to known PfPI4KIIIB inhibitor KDU731, we sequenced the *pfpi4kiiib* gene in our resistant parasites recovering an E1361D mutation which was located near several other *pfpi4kiiib* drug resistance mutations ^37-39,41^. Examination of a homology structure of PfPI4KIIIB indicated the resistance mutations clustered closely to the binding sites of MMV390048 and M-797 predicted by *in silico* docking, providing a rationale for how the E1361D resistance mutation could reduce the affinity of OGHL250-like compounds for the kinase ^39^. The M-797-resistant parasites were refractory to the pyridyl-furan lead compounds **WEHI-518** and **C3** but not to **OGHL250** indicating that during the development of the lead compounds, their specificity for PfPI4KIIIB increased while activity against other proteins targeted by **OGHL250** decreased.

Several inhibitory lead compounds to PfPI4KIIIB have already been developed, some of which have recently undergone phase 1 clinical trials ^46,47^. The treatment of schizonts with the PfPI4KIIIB inhibitor KAI407, closely related to KDU691, has been previously observed to produce poorly invasive merozoites ^39^. This could be due to defective plasma membrane ingression in the schizont, disrupting the proper formation of membranes around the developing merozoites. To investigate this further, we examined Giemsa-stained smears of inhibitor-treated parasites after the 4-hour invasion assay and found many young, malformed rings that failed to grow properly even 20 hours after compound removal. This indicates the pyridyl-furan compounds could also be inhibiting the conversion of newly invaded merozoites into ring-stage parasites.

In contrast to developing merozoites, fully formed merozoites mechanically released from their old RBCs could efficiently invade new RBCs in the presence of KAI407 ^39^. We found this was also the case for M-797 and contrasts with our observations for **OGHL250**, **WEHI-518** and **C3** where purified merozoites appeared to poorly invade RBCs in the presence of these compounds. Based on our observations of Giemsa-stained smears of merozoites that were allowed to egress and invade over a 4-h window, some of the purified merozoites may have invaded but failed to differentiate into rings. It is possible that **OGHL250**, **WEHI-518** and **C3** could block important protein trafficking events essential for RBC invasion such as the secretion of the merozoites’ microneme or rhoptry organelles. It is also likely that the compounds were inhibiting membrane and protein trafficking events required for the differentiation of newly invaded merozoites into rings. If these events depend upon PfPI4KIIIB, it is unknown why KAI407 and M-797 did not inhibit invasion. It is possible that **OGHL250**, **WEHI-518** and **C3** may have targets additional to PfPI4KIIIB which could explain why their EC_50_s for the 4-hour and purified merozoite invasion assays are higher than for growth.

PfPI4KIIIB phosphorylates phosphatidylinositol to produce phosphatidylinositol 4-phosphate. Although phosphoinositides only make up about 1% of membrane phospholipids they play important roles in many cellular processes such as signal transduction, membrane trafficking and protein transport ^48^. Apart from position 4, the inositol head group can be phosphorylated singly and multiply at several other positions by different phosphoinositol kinases to produce up to seven types of phosphoinositol phosphate (PIPs). Through enrichment at different organellar membranes and locations, the PIPs act as a code to regulate membrane-dependent cellular functions. Recent investigation of PIPs in *P. falciparum* using protein reporters that recognise the different PIP types indicated that PI4P is found at the Golgi apparatus and plasma membrane, PI3P concentrates at the apicoplast, plasma membrane and food vacuole membranes and PI5P is found at the plasma membrane, nuclear membrane and possibly the transitional ER in schizonts ^48^. The multiply phosphorylated PIP species PI(3,4)P2 and PI(3,4,5)P3 are cytosolic, and PI(4,5)P2 is in the PM and possibly cytostome, a structure which endocytoses the RBC cytosol. PI(3,5)P2 was not detected, indicating *Plasmodium* may not utilise the full complement of PIP types ^48^. In developing merozoites, PI4P is found in the plasma membrane and an internal punctate structure. In mature, free merozoites, PI4P disappears from the plasma membrane but is retained within an internal structure that does not appear to overlap with the microneme or rhoptry organelles that release proteins involved in invasion ^48^. While the PI4P produced by PfPI4KIIIB appears to play a role in merozoite development, it is unclear what role PI4P might play in invasion. Although our data indicate PfPI4KIIIB is important for successful invasion it is possible that PI4P may not be required for merozoite penetration of the RBC but for downstream events where the internalised merozoite must rapidly differentiate into an amoeboid ring-stage parasite over several minutes ^49^. This would involve disassembling many internal membranous structures, such as the inner membrane complex and apical organelles, as well as the secretion and export of parasite proteins into the RBC compartment to remodel the host cell for subsequent rapid growth, replication, and immune evasion ^50-52^. Metamorphosis of the extracellular merozoite into an intraerythrocytic form that feeds and replicates would require extensive reorganisation of membranous structures and proteins, and this is likely heavily dependent on PI4P and the other PIPs.

In conclusion, we have identified the *P. falciparum*-killing compound **OGHL250**, from OGHL library of Merck KGaA, Darmstadt, Germany that also appears to inhibit merozoite invasion of RBCs. From **OGHL250**, the highly potent analogues **WEHI-518** and **C3** were produced that target PfPI4KIIIB, an enzyme required to produce PI4P, which has an important role in vesicular membrane and protein trafficking from the ER to the plasma membrane via the Golgi. PI4P can also be converted into PI4,5P2 and PI3,4,5P3 which may also have important invasion-related functions ^48^. M-797-resistant parasites containing an E1316D mutation in PfPI4KIIIB are also resistant to **WEHI-518** and **C3** but not to **OGHL250**, which could be targeting another kinase involved in invasion. The E1316D mutation clusters with several other resistance mutations to PfPI4KIIIB inhibitors, some of which are undergoing clinical trials in humans such as MMV390048 ^46^. PfPI4KIIIB is an excellent target with inhibitors blocking multiple parasite lifecycle stages ^38,39^. While **WEHI-518** and **C3** are as potent as other leading PfPI4KIIIB inhibitors, they are metabolically unstable and not suitable for further clinical development. Despite this, our work does shed light on the mechanism of action of this important future drug target.

## MATERIALS AND METHODS

### Compounds

All compounds (zaprinast, heparin, artemisinin, chloroquine, brefeldin A and Torin 2) were sourced from Merck KGaA, Darmstadt, Germany unless otherwise stated. C1 (4-[2-(4-fluorphenyl)-5-(1-methylpiperidine-4-yl)-1H pyrrol-3-yl]pyridine) and KDU731 were made according to published protocols^34,53^. WR99210 was from Jacobus Pharmaceutical Company.

### In vitro culturing of *P. falciparum* and parasite strains

Parasites were cultured in human red blood cells (Australian Red Cross Blood Bank) at 4% hematocrit in RPMI media (Sigma-Aldrich, UK), supplemented with 0.2% NaHCO_3_ (Thermo Scientific, Australia), 0.25% Albumax II (Gibco, New Zealand), 0.37 mM hypoxanthine (Sigma, USA), 25 mM HEPES (Gibco, USA), 31.25 mg/L Gentamicin (Gibco, USA) (cRPMI) and incubated at 37°C with 1% O_2_, 5% CO_2_, 94% N_2_. The parasite lines used in this study includes the 3D7 African wild-type strain parasite transfected with a Nanoluciferase reporter proteins targeted to the RBC compartment (Hyp1-Nluc) ^24^ and Dd2 strain parasite with a mutated exonuclease domain within the DNA polymerase δ (Dd2-Polδ) ^35^. Hyp1-Nluc was cultivated under 2.5 nM WR99210 (Jacobus Pharmaceutical Company) selection to maintain episomal expression of the Hyp1-Nluc gene ^54^.

To produce synchronous parasites for protein export and invasion assays they were first synchronized with 5% sorbitol to enrich for ring-stage parasites ^55^. The parasite were grown until late schizonts and layered over a 67% Percoll density gradient buffer in cRPMI (Percoll, 10 mM NaH_2_PO_4_, 143 mM NaCl (GE Healthcare Bio-Sciences, Sweden)) and centrifuged (1500 g / 15 mins) ^56^. The dark-coloured synchronous schizont layer was removed and used for downstream experiments.

### Screening of the Open Global Health Library for *P. falciparum* growth inhibitors

The assay was conducted on sorbitol-synchronised ring-stage Hyp1-Nluc parasites at 2% hematocrit and 0.3% parasitaemia in a 96-well U-bottom plate (Thermo Scientific, Australia) for 72 h in the presence of 2 µM of the 250 compound OGHL library. The plate was subsequently stored at -80°C and thawed at room temperature for 4 h to lyse parasites. Parasite growth was quantified through colorimetric measurement of parasite lactate dehydrogenase activity (LDH) ^57,58^. To measure parasite Nluc activity, 5 μL of resuspended lysed parasite cultures were dispensed into a 96-well white luminescent plate, prior to the addition of 45 μL of 1 X NanoGlo Lysis buffer to lyse the infected RBCs (iRBCs), releasing Nluc. The lysis buffer contained 1:4000 NanoGlo (furimazine) substrate (Promega, USA), to generate a bioluminescence signal through substrate hydrolysis ^59^. The total Nluc signal was then measured with a CLARIOstar multimode plate reader (BMG Labtech).

### Nluc counter screen

The assay was adapted from ^15^ where Hyp1-Nluc schizonts at 1% hematocrit and 1-2% parasitaemia were lysed in 1x NanoGlo buffer (Promega, USA) within a 96-well flat-bottom plate and parasite lysates were added to compounds and incubated for 10 minutes at 37°C. Nluc activity was measured as above. Nluc activity detected in the presence of compounds was normalised to vehicle control (0.1% DMSO).

### 30-minute egress assay

The assay is adapted from ^15^ and conducted on Hyp1-Nluc schizonts at early-to mid-stages (36-40 hpi) isolated via Percoll purification, at 3-5% parasitaemia at 1% hematocrit in a 96-well U-bottom plate with a 30-minute drug-treatment and incubation period at 37°C. The supernatant was removed for Nluc activity detection as above. The assay included a background control for egress comprising of culture media of Percoll-purified cultures kept at 4°C during the 30-minute incubation period, which accounted for potential Nluc signal from spontaneous lysis or cell leakage. The background control was subtracted from the egress values and normalised relative to the vehicle control (0.1% DMSO).

### 4-hour egress/invasion assay

The assay was performed as per ^15^, with background controls subtracted from both the egress and invasion values and normalised relative to the vehicle control (0.1% DMSO)^15^.

### 72-hour growth inhibition assay with Dd2 parasites and the MMV390048 resistant Dd2 with S743T mutation in PI4KIIIB (Swiss TPH)

The testing was performed with the modified [^3^H]-hypoxanthine incorporation assay, as previously reported ^60^.

### Chemistry

See supplementary Experimental and Chemistry sections.

### In vitro drug resistance selection of *P. falciparum* Dd2-Polδ

Dd2-Polδ mutant parasites were used for *in vitro* drug resistance selection to increase the likelihood of selection of random mutations for drug resistance accelerating the selection time ^35^. Dd2-Polδ mixed-stage parasite cultures at 5% parasitaemia and 4% hematocrit were dispensed into each well of a six-well flat-bottom plate (Thermo Scientific, Australia). Five cultures were treated with 10x the growth EC_50_ of **OGHL250** (9.1 µM) and M-797 (4.8 µM), and one was treated with the vehicle control (DMSO) at the same final volume used in the compound-treated wells. Parasite cultures were exposed to the compound until they began to die, and the drug was removed. The drug treatment was repeated when the parasite population had regrown, and this drug-cycling process was repeated five times. Following this, 72 h growth assays in the presence of inhibitory compounds were performed to determine if the parasites had developed resistance. Clonal parasite lines were derived from resistant populations by limiting dilution and retested for resistance before isolation of genomic DNA.

### Known and putative PfPI4KIIIB inhibitors block protein export in *P. falciparum*

Trophozoite stage 20-24 hours post invasion (hpi), *P. falciparum* parasites expressing a Nluc reporter were treated with PfPI4KIIIB inhibitory compounds for 5 hours before being assayed for inhibition of Nluc reporter secretion and export as previously described ^23^.

### Sequence analysis of PfPI4KIIIB gene sequence from resistant parasite

Genomic DNA was isolated from two clones resistant to M-797and two parental clones. Most of *pfpki4kiiib* was amplified as five overlapping PCR fragments using Phusion DNA polymerase (Thermo) as per manufacturer’s instruction. Primer sequences are listed in Table S5 and as only PCR fragment 4 was found to contain a mutation in the resistant parasites, two clones each from another three resistant populations were ligated into the pJet 1.2 plasmid (Thermo) and were sequenced. Sequence data was aligned an analysed in SnapGene.

### Invasion inhibition assay with purified merozoites

Merozoites were grown and purified, inhibitor treated and assayed for invasion as described in ^15^.

## AUTHOR CONTRIBUTIONS

DL, OL, WN, BS and PG conceived this research and designed the experiments. DL, OL, and WN conducted the experiments and generated the figures. PG wrote the manuscript. All authors edited the manuscript. BS, BC and PG provided funding. All authors have approved the submitted manuscript.

## ACKNOWLEDGEMENTS

We are grateful to Merck KGaA Darmstadt, Germany for provison of the Open Global Helath library and to Sven Lindemann for supplying additional compound material. We thank Lifeblood Biological Resources for providing blood and Medicine for Malaria Venture for faciltating the testing of three compounds in the hypoxanthine incorporation assay.

## FUNDING

This work was supported by funding from the Victorian Operational Infrastructure Support Program received by the Burnet Institute. Funding was provided by the National Health and Medical Research Council (grant numbers 1185354, 2001073 and 119780521).

## FIGURES

## Notes

### Competing Interest Statement

The authors have declared no competing interest.

