## Supplementary figures for "A pyridyl-furan series developed from Open Global Health Library blocks red blood cell invasion and protein trafficking in *Plasmodium falciparum* through potential inhibition of the parasite’s PI4KIIIb enzyme"

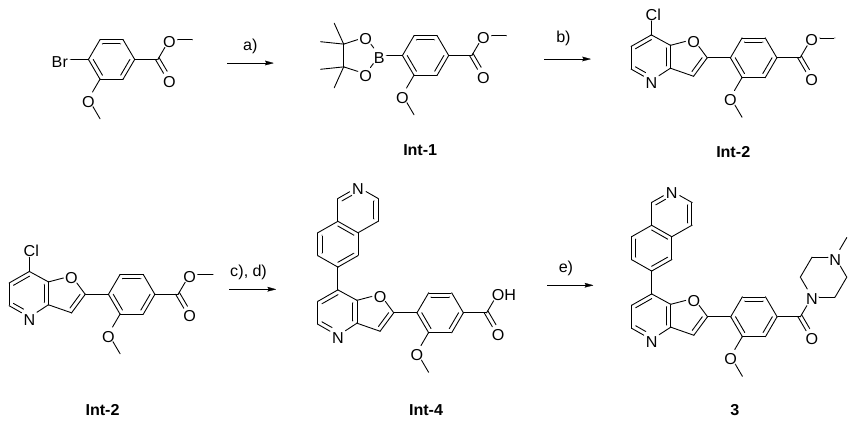

Scheme S1. Synthetic route to generate **3**. *Reagents and conditions:* (a) bis(pinacolato)diboron, potassium acetate, Pd(dppf)Cl_2_.CH_2_Cl_2_, 1,4-dioxane, 80 °C; (b) 7-chloro-2-iodo-furo[3,2-b]pyridine, K_2_CO_3_, Pd(dppf)Cl_2_.CH_2_Cl_2_, 1,4-dioxane/H_2_O (5:1), 100 °C; (c) 6-isoquinolylboronic acid, SPhos, K_2_CO_3_, Pd(OAc)₂ 1,4-dioxane/H_2_O (5:1), 100 °C; (d) LiOH, 1,4-dioxane/H_2_O (5:1); (e) 1-methylpiperazine, HATU, ACN, 20 °C.

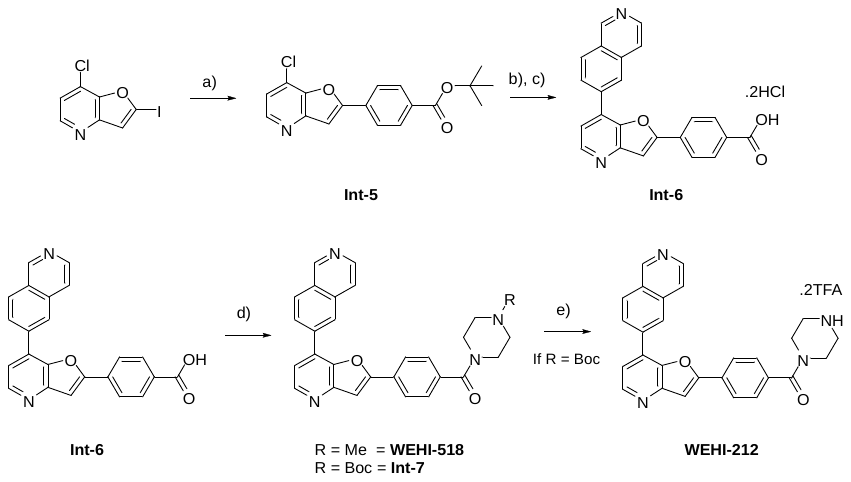

Scheme S2. Synthetic route to generate **WEHI-518** and **WEHI-212**. *Reagents and conditions:* (a) (4-*tert*-butoxycarbonylphenyl)boronic acid, K_2_CO_3_, Pd(dppf)Cl_2_.CH_2_Cl_2_, 1,4-dioxane/H_2_O (5:1), 100 °C; (b) 6-isoquinolylboronic acid, SPhos, K_2_CO_3_, Pd(OAc)₂ 1,4-dioxane/H_2_O (5:1), 100 °C; (c) 4M HCl in 1,4-dioxane; (d) 1-methylpiperazine or Boc-piperazine, HATU, ACN, 20 °C; (e) DCM/TFA (3:1), 20 °C.

### Table S2. Aqueous solubility and *in vitro* metabolism for WEHI-518 and WEHI-212.

| **Cmpd** | **Aqueous Solubility** | | **Mouse Liver Microsomes** | |
| --- | --- | --- | --- | --- |
|  | pH 6.5 (µM) ^a^ | pH 2.0 (µM) ^a^ | *in vitro* CL_int_ (µL/min/mg) | Half-life (min) |
| **WEHI-518** | 160-320 | >320 | 301 | 4.6 |
| **WEHI-212** | 20-40 | 160-320 | 180 | 7.7 |

^a^ Kineitc solubility estimated by nephelometry.

### Table S3. Human kinase activity of WEHI-518 in ATP-Glo assays.^a^

| **Cmpd** | human kinase activity IC_50_ (µM) | | | |
| --- | --- | --- | --- | --- |
|  | PI4Ka | IKKe | Syk | TBK |
| **WEHI-518** | 3.0 | 0.22 | 0.27 | >10 |

^a^ Data generated by Reaction Biology.

Table S4. 72-hour growth assay EC_50_ of compounds against Dd2 parasites and the MMV390048 resistant Dd2 with S743T mutation in PI4KIIIB.

| **Cmpd** | **Parasite line EC_50_ (SD)*** | |
| --- | --- | --- |
|  | **Dd2** | **Dd2 (PI4KIIIB S743T)** |
| **MMV396797** | 0.295 (0.044) | 0.544 (0.036) |
| **OGHL250** | 0.221 (0.016) | 0.192 (0.056) |
| **WEHI-518** | 0.005 (0.0002) | 0.022 (0.0005) |

* N=2-3 biological replicates.

### Table S5. PCR primers used to amplify *pfpi4kiiib* gene for sequencing

| PI4K.1F | ATCATGGTTATATCATTCCATTGTTGAAGATAATTCTAGT |
| --- | --- |
| PI4K.1R | TACTTTCTAATGGTATACACAAACCTGTCATGGCA |
| PI4K.2F | TGAATATGAGAAGATGTATAGTGGCTTGTTGTGA |
| PI4K.2R | AGATGATGAAGATAAATTATCATTTGCTGTTGGAACT |
| PI4K.3F | TGAAAAGAAAAACTTTGAAGAAGAACAAAAGGGA |
| PI4k.3R | AGATCCCACGTTTTTAATTTTCCATAAGGAGA |
| PI4K.4F | ATAACCAAGATAATACCATTAGTAACTTCCCAAACA |
| PI4K.4R | TCTGAATGTTTTCTAGCCTCTAGAAAGCCACT |
| PI4K.5F | TGTGAATTTTGAAACATCCCCATTCAAATTAACACA |
| PI4K.5R | TCACATAATTCCATTTGTTATTCTTTGAAAGTAGTCA |

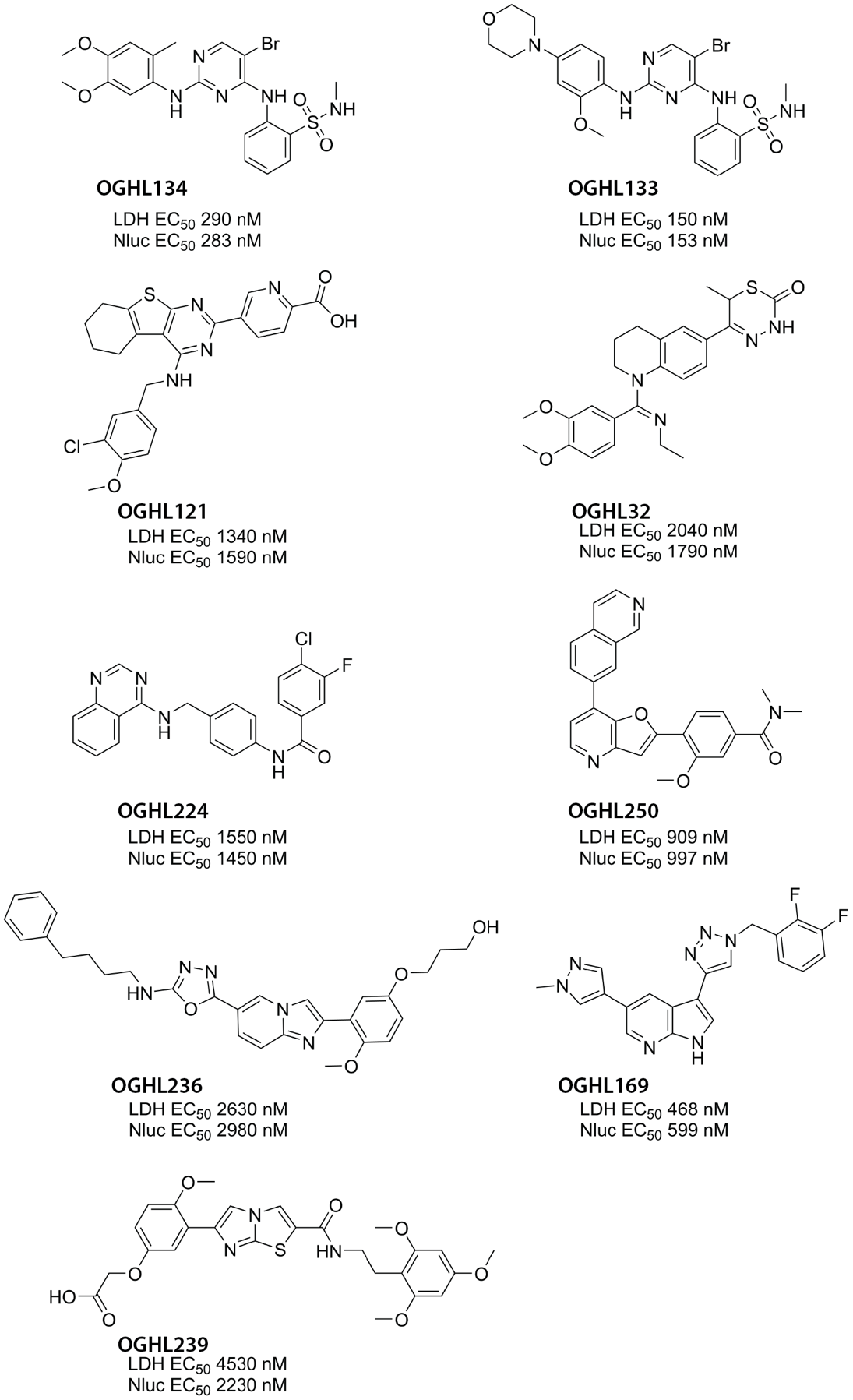

Supplementary Figure 1. Structures of *Plasmodium falciparum* growth inhibitory compounds from OGHL compound library. Growth assays were performed on transgenic *P. falciparum* 3D7 strain parasites expressing an exported nanoluciferase (Nluc) reporter for 72 hours in the presence of serially diluted compounds. EC_50_ values for lactate dehydrogenase (LDH) and Nluc activities are indicated. * Subsequent counter screen indicates OGHL239 can inhibit Nluc activity.

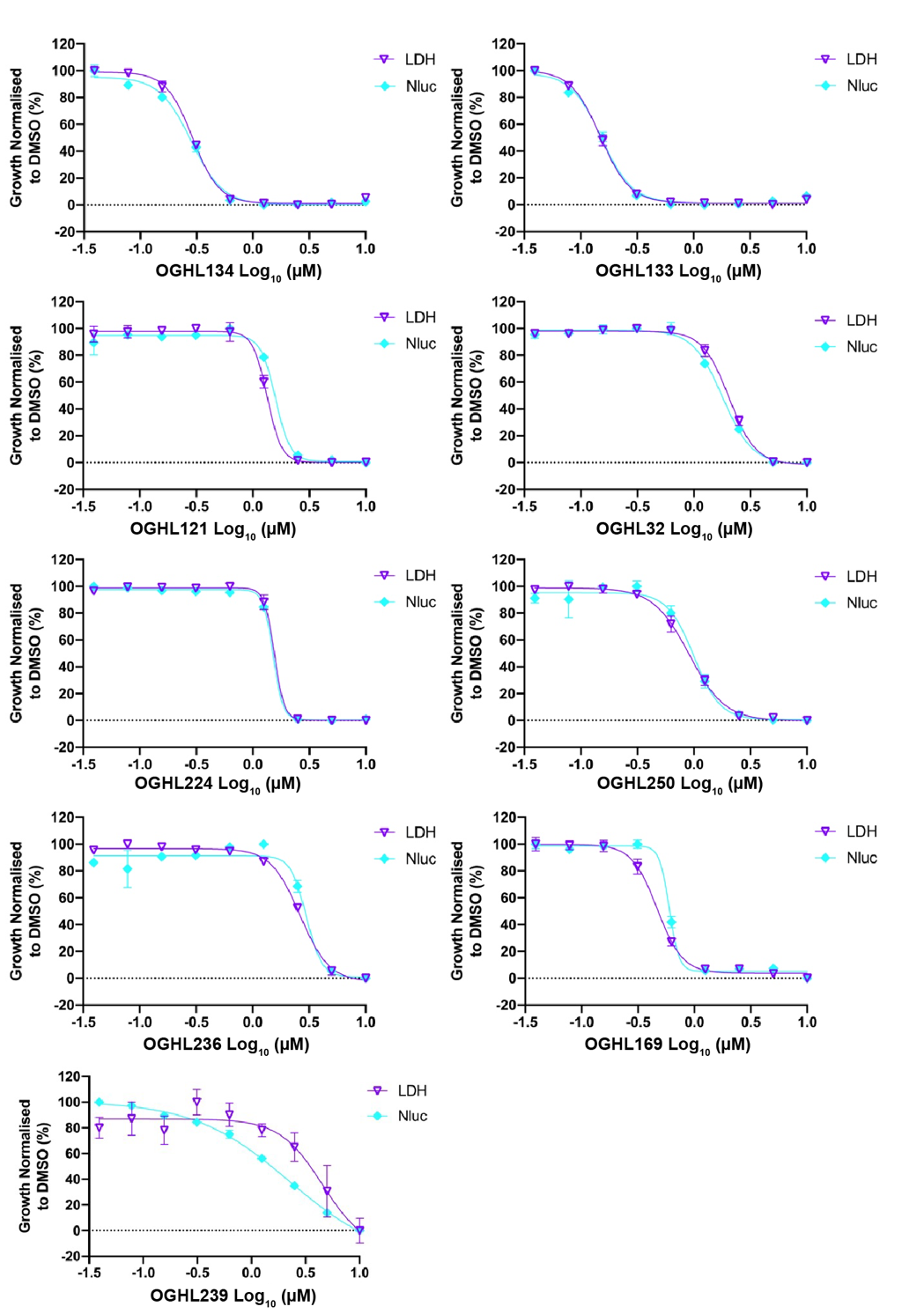

Supplementary Figure 2. Dose-response curves and growth EC_50_ values of Open Global Health Library hit compounds against parasite growth validates their growth inhibitory activity, except for OGHL239 (likely a direct Nanoluciferase (Nluc) inhibitor). To determine their potencies (parasite growth EC_50_), the 72-hour growth assay on the OGHL hit compounds serially diluted one in two over nine concentrations starting from 10 µM was conducted. Except for OGHL239, the similarity in the growth EC50 values detected for the hit compounds via lactate dehydrogenase (LDH) and Nluc activity confirms their parasite growth inhibition activity. Given the relatively large 95% confidence interval of the EC50 values (C.I.) of OGHL239 using LDH detection compared to Nluc detection, OGHL239 is likely a direct Nluc inhibitor.

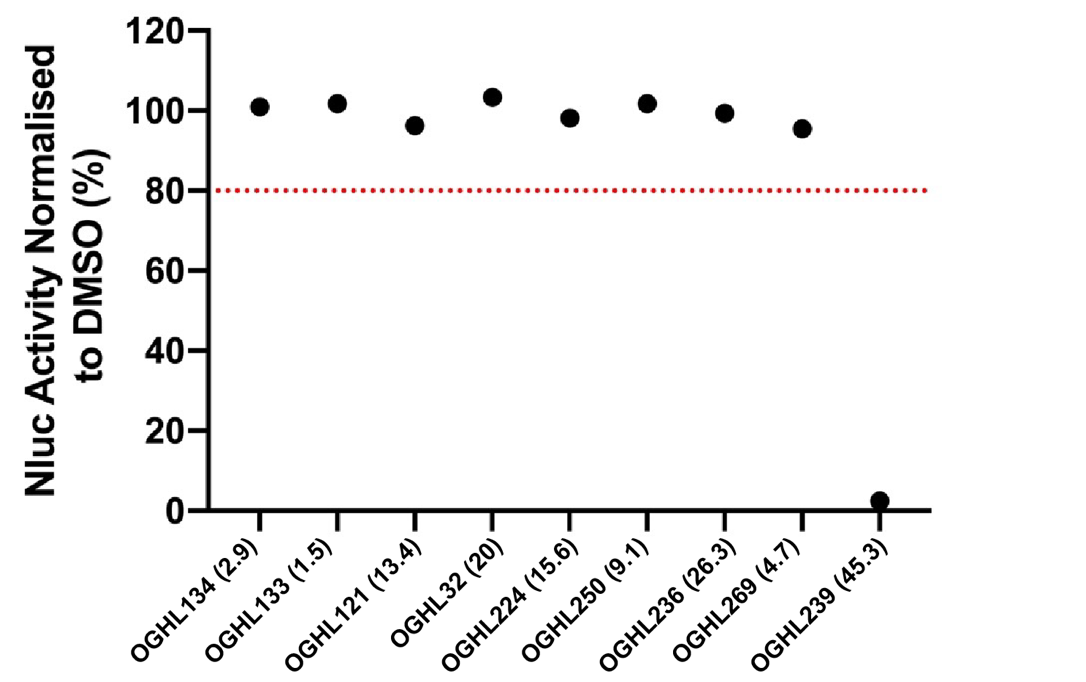

Supplementary Figure 3. Nanoluciferase (Nluc) activity counter screen indicates that OGHL239 is a direct Nluc inhibitor. The counter screen measured the direct activity of the Open Global Health Library (OGHL) hit compounds against Nluc. The screen demonstrates that OGHL239 is direct Nluc inhibitor. All values were normalised to vehicle control (0.1% DMSO) and each dot represents the mean of a compound from three technical replicates. Dotted line indicates cut-off value of 80% for Nluc activity. OGHL compounds were tested at 10x EC_50_ of growth, all concentrations specified in brackets are in μM.

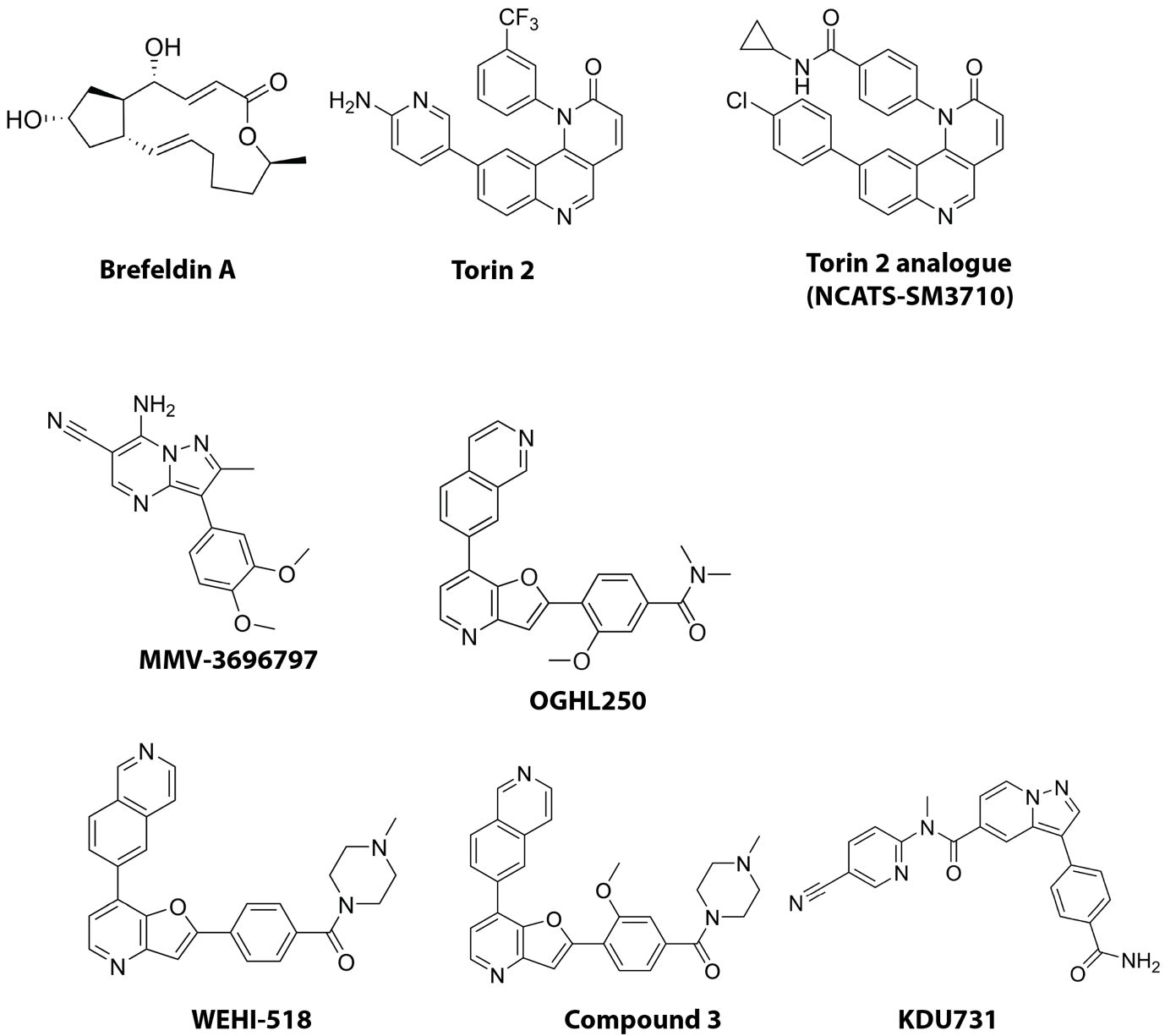

Supplementary Figure 4. Structures of compounds used for protein secretion and export screen.

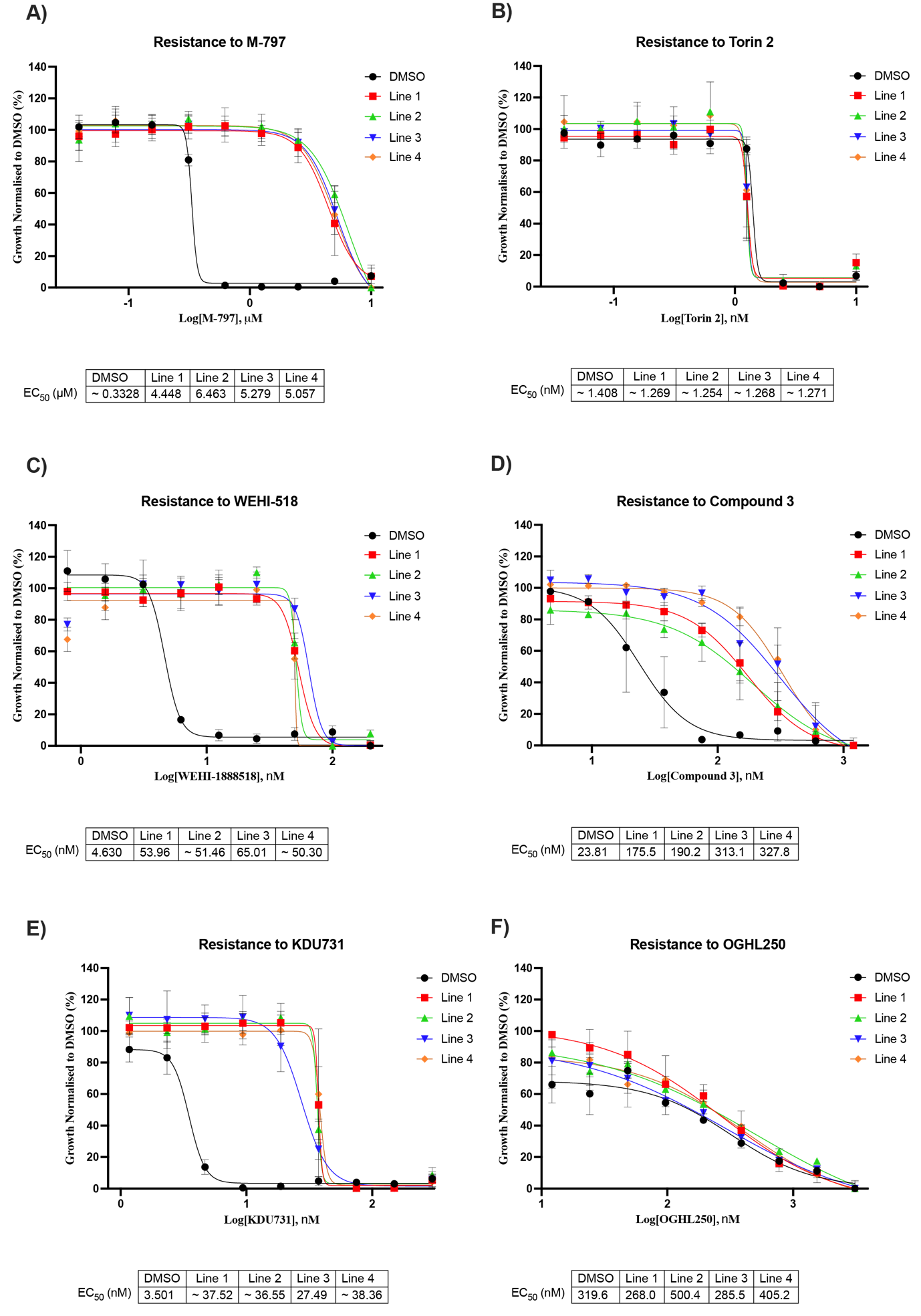

Supplementary Figure 5. Growth of MMV396797 resistant parasite lines on known and putative PfPI4KIIIB inhibitors. (A-F) Four resistant parasite lines and the DMSO treated Parental parasites were grown for 74 hours on a nine-point compound dilution series as indicated. Parasite growth was evaluated by measuring lactate dehydrogenase activity. EC_50_ in µM or nM for the compounds are indicated.

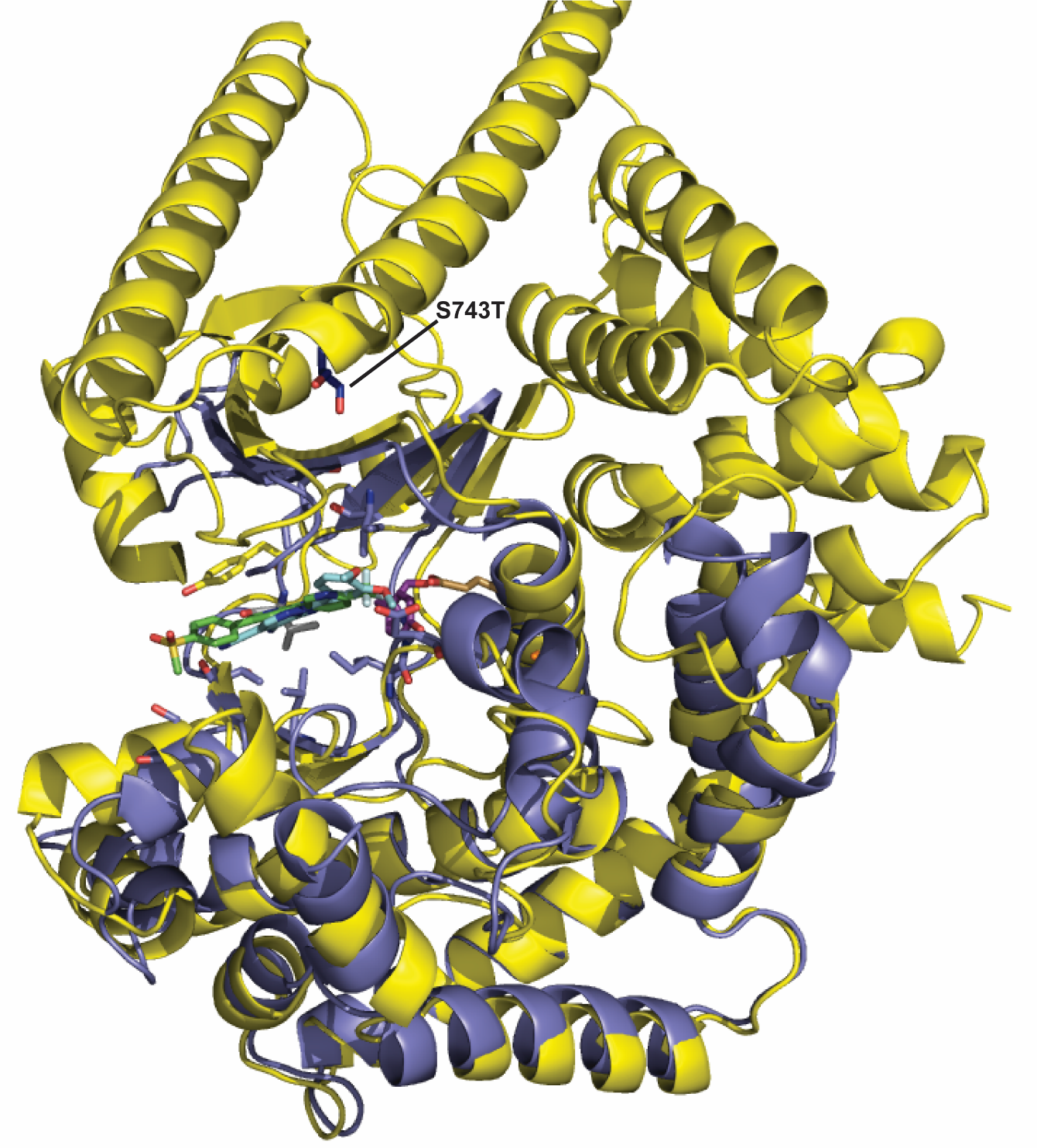

**Figure S6.** ALPHA-fold (Jumper et al 2021, Vardi et al. 2022) model of Pf PI4K (PF3D7_0509800) in yellow overlaid with the homology model of Pf PI4K shown in Figure 8B. The ALPHA-fold model shows structural domains other than the kinase domain highlighting the proximity of S743 (in dark blue) which is the amino acid that is mutated to Thr in MMV390048 resistant parasites. Amino acids 1-269, 502-709 and 842-987 are truncated in the ALPHA-fold model for clarity.

[Jumper, J *et al*. Highly accurate protein structure prediction with AlphaFold. *Nature* (2021).](https://www.nature.com/articles/s41586-021-03819-2)

[Varadi, M *et al*. AlphaFold Protein Structure Database: massively expanding the structural coverage of protein-sequence space with high-accuracy models. *Nucleic Acids Research* (2022).](https://academic.oup.com/nar/advance-article/doi/10.1093/nar/gkab1061/6430488)

### EXPERIMENTAL SECTION

**General Chemistry Methods.** NMR spectra were recorded on a Bruker Ascend^TM^ 300. Chemical shifts are reported in ppm on the δ scale and referenced to the appropriate solvent peak. MeOD, DMSO-*d_6_*, D_2_O, and CDCl_3_ contain H_2_O. Chromatography was performed with silica gel 60 (particle size 0.040-0.063 µm) using an automated CombiFlash Rf Purification System. LCMS were recorded on an Agilent LCMS system comprised of an Agilent G6120B Mass Detector, 1260 Infinity G1312B Binary pump, 1260 Infinity G1367E HiPALS autosampler, and 1260 Infinity G4212B Diode Array Detector (Method B). Conditions for LCMS Method A were as follows, column: Luna® Omega 3 µm PS C18 100 Å, LC Column 50 × 2.1 mm at 20 ºC, injection volume 2 µL, gradient: 5-100% B over 3 min (solvent A: H_2_O 0.1% formic acid; solvent B: ACN 0.1% formic acid), flow rate: 1.5 mL/min, detection: 100-600 nm, acquisition time: 4.3 min. Conditions for LCMS Method B were as follows, column: Poroshell 120 EC-C18, 2.1 × 30 mm 2.7 Micron at 30 ºC, injection volume 2 µL, gradient: 5-100% B over 3 min (solvent A: H_2_O 0.1% formic acid; solvent B: ACN 0.1% formic acid), flow rate: 0.8 mL/min, detection: 254 nm, acquisition time: 4.1 min. Unless otherwise noted, all compounds were found to be >95% pure by this method. Compounds **OGHL250**, **1** and **2** were procured from the Merck OGHL library (https://www.merckgroup.com/en/research/open-innovation/biopharma-open-innovation-portal/open-global-health-library.html.) and used without further purification.

### Chemistry Procedures.

**Int-1**

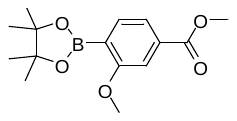

*Methyl 3-methoxy-4-(4,4,5,5-tetramethyl-1,3,2-dioxaborolan-2-yl)benzoate* (**Int-1**). To a solution of methyl 4-bromo-3-methoxy-benzoate (2.00 g, 8.16 mmol) in 1,4-dioxane (25 mL) were added bis(pinacolato)diboron (2490 mg, 9.79 mmol), Pd(dppf)Cl_2_.CH_2_Cl_2_ (330 mg, 0.41 mmol) and potassium acetate (2.45 g, 24.5 mmol) at room temperature. The resulting mixture was stirred overnight at 80 °C. The reaction mixture was cooled to 20 ℃ and quenched by the addition of water (10 ml). The resulting mixture was extracted with ethyl acetate (3 × 20 mL). The organic phases were combined, washed with brine (20 mL), dried over anhydrous Na_2_SO_4_ and concentrated. The crude was then purified by column chromatography eluting with 100% heptane to 50% EtOAc to afford **Int-1** as an oil (1.0 g, 44%). ^1^H NMR (300 MHz, CDCl_3_): δ 7.71 (d, *J* 7.6 Hz, 1H), 7.61 (dd, *J* 7.58, 1.30 Hz, 1H), 7.49 - 7.53 (m, 1H), 3.93 (s, 3H), 3.90 (s, 3H), 1.37 (s, 12H), 1.27 (s, 6H). LCMS m/z 293.2 [M+1].

**Int-2**

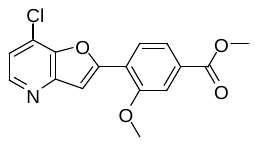

*Methyl 4-(7-chlorofuro[3,2-b]pyridin-2-yl)-3-methoxy-benzoate* (**Int-2**). To a solution of **Int-1** (1.0 g, 3.4 mmol) in 1,4-dioxane (20 mL) and H_2_O (4 mL) was added 7-chloro-2-iodo-furo[3,2-b]pyridine (0.96 g, 3.4 mmol), K_2_CO_3_ (1.13 g, 8.19 mmol) and Pd(dppf)Cl_2_.CH_2_Cl_2_ (270 mg, 0.33 mmol) at 20 ℃. The resulting mixture was stirred for 3 h at 100 ℃. The reaction mixture was cooled to 20 ℃ and quenched by the addition of H_2_O (10 ml). The mixture was then extracted with EtOAc (3 × 15 mL). the organic phases were combined, washed with brine (20 mL) dried over anhydrous Na_2_SO_4_ and concentrated. The crude was then purified by column chromatography eluting with 100% heptane to 100% EtOAc to afford **Int-2** (660 mg, 61%). ^1^H NMR (300 MHz, CDCl_3_): δ 8.45 (br s, 1H), 8.21 (d, *J* 8.2 Hz, 1H), 7.80 (dd, *J* 8.2, 1.4 Hz, 1H), 7.67 - 7.75 (m, 2H), 7.29 (d, *J* 4.7 Hz, 1H), 4.10 (s, 3H), 3.98 (s, H). LCMS m/z 318.2 [M+1].

**Int-3**

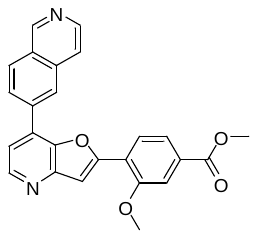

*Methyl 4-[7-(6-isoquinolyl)furo[3,2-b]pyridin-2-yl]-3-methoxy-benzoate* (**Int-3**). To a solution of **Int-2** (300 mg, 0.94 mmol) in 1,4-dioxane (5 mL) and H_2_O (1 mL) was added 6-isoquinolylboronic acid (240 mg, 1.4 mmol), SPhos (116 mg, 0.283 mmol) K_2_CO_3_ (390 mg, 2.8 mmol), Pd(OAc)₂ (21 mg, 0.094 mmol) at 20 ℃. The vial was purged with nitrogen gas for 5 min and then stirred for 6 h at 100 ℃. This was then cooled to 20 ℃ and treated with H_2_O (5 ml). The resulting solution was extracted with DCM (3 × 10 mL). The organic phases were combined, washed with brine (20 mL), dried over anhydrous Na_2_SO_4_ and concentrated. The crude was then purified by column chromatography eluting with 100% DCM to 10% MeOH to afford **Int-3** as a solid (110 mg, 29%). ^1^H NMR (300 MHz, CDCl_3_): δ 9.49 (s, 1H), 8.74 (d, *J* 5.3 Hz, 1H), 8.69 (d, J 6.3 Hz, 1H), 8.61 (s, 1H), 8.38 (s, 2H), 8.11 (d, *J* 8.2 Hz, 1 H), 8.03 (d, *J* 5.21 Hz, 1H), 7.89 (s, 1H), 7.82 (dd, *J* 8.1, 1.3 Hz, 1 H), 7.77 (s, 1 H), 7.62 (d, *J* 6.1 Hz, 1 H), 4.16 (s, 3H), 4.00 (s, 3H). LCMS m/z 411.2 [M+1].

**Int-4**

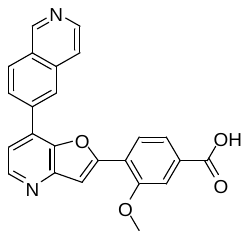

*4-[7-(6-Isoquinolyl)furo[3,2-b]pyridin-2-yl]-3-methoxy-benzoic acid* (**Int-4**). **Int-3** (80 mg, 0.19 mmol) was dissolved in a 1:1 mixture of MeOH and H_2_O (10 mL). LiOH (23.3 mg, 0.975 mmol) was added, and the reaction stirred at 50 °C for 1 h. The MeOH was then removed in vacuo and the reaction neutralized with citric acid (pH 6). The resulting precipitate was then filtered and dried under vacuum to afford **Int-4** as a white solid (77 mg, 100%). ^1^H NMR (300 MHz, DMSO-d_6_) δ 9.40 (s, 1H) 9.45 (s, 1H) 8.73 (s, 1H) 8.65 – 8.62 (m, 1H) 8.58 (d, *J* 5.7 Hz, 1H), 8.49 (s, 1H), 8.41 (s, 1H), 8.29 (s, 1H), 8.22 (d, J 1.7 Hz, 1H), 7.95 (d, *J* 7.7 Hz, 2H), 7.75 (d, *J* 5.1 Hz, 1H), 7.70 (s, 1H), 7.64 – 7.62 (m, 1H) 4.04 (s, 3H). LCMS m/z 397.2 [M+1].

**3**

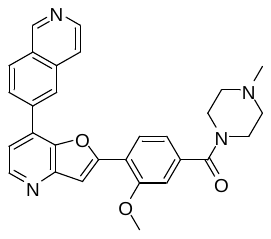

*[4-[7-(6-Isoquinolyl)furo[3,2-b]pyridin-2-yl]-3-methoxy-phenyl]-(4-methylpiperazin-1-yl)methanone* (**3**)**. Int-4** (15 mg, 0.038 mmol), HATU (21.6 mg, 0.0568 mmol) and 1-methylpiperazine (0.042 mL, 0.38 mmol) were stirred ACN (1 mL) for 16 h. The reaction was then quenched with 5% citric acid and concentrated. The crude was then dissolved in EtOAc (10 mL) and washed with saturated NaHCO_3_ (10 mL), brine (10 mL), dried over anhydrous Na_2_SO_4_ and concentrated. The crude was then purified by column chromatography eluting with 100% DCM to 20% MeOH/ammonia solution (10:1) to afford **3** (3.90 mg, 22%). ^1^H NMR (300 MHz, CDCl3): δ 9.38 (s, 1H), 8.67 (dd, *J* 7.3, 5.4 Hz, 2H), 8.48 (s, 1H), 8.18 - 8.28 (m, 2H), 8.06 (d, *J* 8.0 Hz, 1H), 7.82 (d, *J* 5.8 Hz, 1H), 7.71 (s, 1H), 7.51 (d, *J* 5.0 Hz, 1H), 7.14 - 7.18 (m, 1H), 7.11 (dd, *J* 7.9, 1.4 Hz, 1H), 4.09 (s, 3H), 3.83 (app s, 2H), 3.52 (app s, 2H), 2.33 - 2.58 (m, 7H). LCMS m/z 479.2 [M+1].

**Int-5**

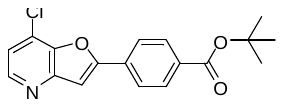

*tert-Butyl 4-(7-chlorofuro[3,2-b]pyridin-2-yl)benzoate* (**Int-5**). The procedure used for **Int-2** was followed using (4-tert-butoxycarbonylphenyl)boronic acid (240 mg, 0.72 mmol) and 7-chloro-2-iodo-furo[3,2-b]pyridine (200 mg, 0.72 mmol) to obtain **Int-5** as a solid (99 mg, 29%). ^1^H NMR (300 MHz, CDCl_3_): δ 8.58 (d, *J* 5.5 Hz, 1H), 8.10 - 8.16 (m, 2H), 8.00 - 8.04 (m, 2H), 7.54 (s, 1H), 7.36 (d, *J* 5.5 Hz, 1H), 3.71 (s, 3H), 1.65 (s, 9H). LCMS m/z 330.2 [M+1].

**Int-6**

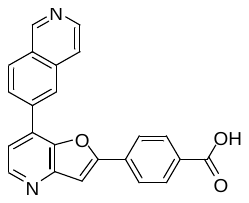

*4-[7-(6-Isoquinolyl)furo[3,2-b]pyridin-2-yl]benzoic acid dihydrochloride* (**Int-6**). The procedure used for **Int-3** was followed using **Int-5** (99 mg, 0.21 mmol) and 6-isoquinolylboronic acid (109 mg, 0.630 mmol) to afford a protected intermediate of **Int-6** as a solid (44 mg, 50 %). This was then dissolved in 4M HCl in dioxane (2 mL) and stirred at 50 ºC for 2 h. The reaction was then concentrated to afford **Int-6** as a solid (46 mg, 50%). ^1^H NMR (300 MHz, CDCl_3_): δ 9.45 (br s, 1H), 8.71 (br s, 2H), 8.52 (br s, 1H), 8.27 (br s, 2H), 8.13 (d, *J* 8.2 Hz, 2H), 7.98 (d, *J* 7.9 Hz, 3H), 7.90 (br s, 1H), 7.56 (br s, 1H), 7.46 (s, 2H). LCMS m/z 367.2 [M+1].

**WEHI-518**

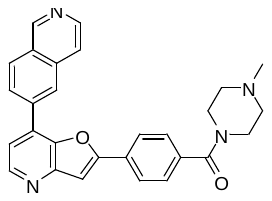

*(4-(7-(isoquinolin-6-yl)furo[3,2-b]pyridin-2-yl)phenyl)(piperazin-1-yl)methanone* (**WEHI-518**). The procedure used for **3** was followed using **Int-6** (15 mg, 0.034 mmol) to afford **WEHI-518** as a solid (8.6 mg, 56%). ^1^H NMR (300 MHz, MeOD): δ 9.39 (s, 1H), 8.72 (s, 1H), 8.65, (d, *J* 5.1 Hz, 1H), 8.58 (d, *J* 5.9 Hz, 1 H), 8.40 (d, *J* 2.7 Hz, 2H), 8.15 (d, *J* 8.3 Hz, 2H), 8.06 (d, J 5.74 Hz, 1H), 7.79 (d, J 5.1 Hz, 1 H), 7.59 - 7.67 (m, 3H), 3.82 (app s, 2H), 3.56 (app s, 2 H), 2.56 (app s, 2 H), 2.49 (app s, 2 H), 2.36 (s, 3H). LCMS m/z 449.2 [M+1].

**WEHI-212**

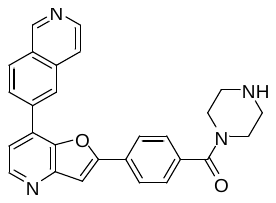

*[4-[7-(6-Isoquinolyl)furo[3,2-b]pyridin-2-yl]phenyl]-piperazin-1-yl-methanone bis trifluoroacetic acid salt* (**WEHI-212**). The procedure used for **3** was followed using **Int-6** (5 mg, 0.011 mmol), 1-Boc-piperazine (4.2 mg, 0.0228 mmol) and DIPEA (5 µL, 0.034 mmol) to afford a protected intermediate (**Int-7**) as a solid (5.1 mg, 84%). This was then dissolved in a mixture of DCM/TFA (3:1, 1 mL) and stirred for 30 min. The reaction was then concentrated and lyophilized to afford **WEHI-212** (6.0 mg, 80%). ^1^H NMR (300 MHz, DMSO-d_6_) δ 9.59 (s, 1H), 8.97 (br s, 2H), 8.83 (s, 1H), 8.68 (d, *J* 5.8 Hz, 1H), 8.73 (d, *J* 5.0 Hz, 1H), 8.48 (s, 2H), 8.10 - 8.27 (m, 3H), 7.95 (s, 1H), 7.84 (d, J 5.1 Hz, 1H), 7.68 (d, *J* 8.4 Hz, 2H), 3.70 (app s, 4 H), 3.21 (app s, 4H). LCMS m/z 435.2 [M+1].
